# CellSAM: A Foundation Model for Cell Segmentation

**DOI:** 10.1101/2023.11.17.567630

**Authors:** Uriah Israel, Markus Marks, Rohit Dilip, Qilin Li, Changhua Yu, Emily Laubscher, Ahamed Iqbal, Elora Pradhan, Ada Ates, Martin Abt, Caitlin Brown, Edward Pao, Shenyi Li, Alexander Pearson-Goulart, Pietro Perona, Georgia Gkioxari, Ross Barnowski, Yisong Yue, David Van Valen

**Author notes:** Contributing authors; sli5@caltech. These authors contributed equally to this work (Uriah Israel, Markus Marks). These authors contributed equally to this work (Rohit Dilip, Qilin Li).

## Abstract

Cells are a fundamental unit of biological organization, and identifying them in imaging data – cell segmentation – is a critical task for various cellular imaging experiments. While deep learning methods have led to substantial progress on this problem, most models are specialist models that work well for specific domains but cannot be applied across domains or scale well with large amounts of data. In this work, we present CellSAM, a universal model for cell segmentation that generalizes across diverse cellular imaging data. CellSAM builds on top of the Segment Anything Model (SAM) by developing a prompt engineering approach for mask generation. We train an object detector, CellFinder, to automatically detect cells and prompt SAM to generate segmentations. We show that this approach allows a single model to achieve human-level performance for segmenting images of mammalian cells, yeast, and bacteria collected across various imaging modalities. We show that CellSAM has strong zero-shot performance and can be improved with a few examples via few-shot learning. Additionally, we demonstrate how CellSAM can be applied across diverse bioimage analysis workflows. A deployed version of CellSAM is available at https://cellsam.deepcell.org/.

## 1 Introduction

Accurate cell segmentation is crucial for quantitative analysis and interpretation of various cellular imaging experiments. Modern spatial genomics assays can produce data on the location and abundance of 10^2^ protein species and 10^3^ RNA species simultaneously in living and fixed tissues ^1–5^. Accurate cell segmentation allows this type of data to be converted into interpretable tissue maps of protein localization and transcript abundances; these maps provide important insights into the biology of healthy and diseased tissues. Similarly, live-cell imaging provides insight into dynamic phenomena in bacterial and mammalian cell biology. Studying live-cell imaging data has provided mechanistic insights into critical phenomena such as the mechanical behavior of the bacterial cell wall ^6,7^, information transmission in cell signaling pathways ^8–13^, heterogeneity in immune cell behavior during immunotherapy ^14^, and the morphodynamics of development ^15^. Cell segmentation is also a key challenge for these experiments, as cells must be segmented and tracked to create temporally consistent records of cell behavior that can be queried at scale. These methods have seen use in several systems, including mammalian cells in cell culture ^13,16^ and tissues ^5^, bacterial cells ^17–20^, and yeast ^21–23^.

Significant progress has been made in recent years on the problem of cell segmentation, driven primarily by advances in deep learning ^24^. Progress in this space has occurred mainly in two distinct directions. The first direction seeks to find deep learning architectures that achieve state-of-the-art performance on cellular imaging tasks. These methods have historically focused on a particular imaging modality (e.g., brightfield imaging) or target (e.g., mammalian tissue) and have difficulty generalizing beyond their intended domain ^25–31^. For example, Mesmer’s ^28^ representation for a cell (cell centroid and boundary) enables good performance in tissue images but would be a poor choice for elongated bacterial cells. Similar trade-offs in representations exist for the current collection of Cellpose models, necessitating the creation of a model zoo ^26,32^. The second direction is to work on improving labeling methodology. Cell segmentation is an application of the instance segmentation problem, which requires pixel-level labels for every object in an image. Creating these labels can be expensive (10^−2^ USD/label, with hundreds to thousands of labels per image) ^28,33^, which provides an incentive to reduce the marginal cost of labeling. A recent improvement to labeling methodology has been human-in-the-loop labeling, where labelers correct model errors rather than produce labels from scratch ^26,28,34^. Further reductions in labeling costs can increase the amount of labeled imaging data by orders of magnitude.

Recent work in machine learning on foundation models holds promise for providing a complete solution. Foundation models are large deep neural network models (typically transformers ^35^) trained on large amounts of data in a self-supervised fashion with supervised fine-tuning on one or several tasks ^36^. Foundation models include the GPT ^37,38^ family of models, which have proven transformative for natural language processing ^36^. These types of attention-based models have recently been used for processing biological sequences ^39–43^. These successes have inspired similar efforts in computer vision. The Vision Transformer (ViT) ^44^ was introduced in 2020 and has since been used as the basis architecture for a collection of vision foundation models ^45–49^. A key feature of foundation models is the scaling of model performance with model size, dataset size, and compute ^50^; these scaling laws have been observed for both language and vision models ^51^. These scaling laws offer a path toward general models for cellular image analysis by increasing dataset and model size in exchange for dealing with the increased compute cost of training foundation models. This is in contrast to prior efforts that have focused on model architecture design and representation engineering.

One recent foundation model well suited to cellular image analysis is the Segment Anything Model (SAM) ^52^. This model uses a ViT to extract information-rich features from raw images. These features are then directed to a module that generates instance masks based on user-provided prompts, which can be either spatial (e.g., an object centroid or bounding box) or semantic (e.g., an object’s visual description). Notably, the promptable nature of SAM enabled scalable dataset construction, as preliminary versions of SAM allowed labelers to generate accurate instance masks with one to two clicks. The final version of SAM was trained on a dataset of 11 million images containing over 1 billion masks and demonstrated strong performance on various zero-shot evaluation tasks. Recent work has attempted to apply SAM to problems in biological and medical imaging, including medical image segmentation ^53–55^, lesion detection in dermatological images ^56,57^, nuclear segmentation in H&E images ^58,59^, and cellular image data for use in the Napari software package ^60^. While promising, these studies reported challenges adapting SAM to these new use cases ^53,60^. These challenges include reduced performance and uncertain boundaries when transitioning from natural to medical images. Cellular images contain additional complications – they can involve different imaging modalities (e.g., phase microscopy vs. fluorescence microscopy), thousands of objects in a field of view (as opposed to dozens in a natural image), and uncertain and noisy boundaries (artifacts of projecting 3D objects into a 2D plane) ^60^.

In addition to these challenges, SAM’s default strategy for automatic prompting does not allow for accurate inference on cellular images. SAM’s automated prompting uses a uniform grid of points to generate masks, an approach poorly suited to cellular images given the wide variation of cell densities. More precise prompting (e.g., a bounding box or mask) requires prior knowledge of cell locations. Because cellular images often contain a large number of cells, it is impractical for users to provide prompts to SAM manually. This limitation makes it challenging for SAM to serve as a foundation model for cell segmentation since it still requires substantial human input for inference. A solution that enables the automatic generation of prompts would enable SAM-like models to serve as foundation models and knowledge engines, as they could accelerate the generation of labeled data, learn from them, and make that knowledge accessible to life scientists via inference.

In this work, we developed CellSAM, a foundation model for cell segmentation (Fig. 1). CellSAM extends the SAM methodology to perform automated cellular instance segmentation. To achieve this, we first assembled a comprehensive dataset for cell segmentation spanning five broad data archetypes: tissue, cell culture, yeast, H&E, and bacteria. Critically, we removed data leaks between training and testing data splits to ensure an accurate assessment of model performance. To automate inference with CellSAM, we developed CellFinder, a transformer-based object detector that uses the Anchor DETR framework ^61^. CellSAM and CellFinder share SAM’s ViT backbone for feature extraction; these features are first used by CellFinder to generate bounding boxes around the cells to be used as prompts for SAM. The bounding boxes (prompts) and ViT features are fed into a decoder to generate instance segmentations of the cells in an image. We trained CellSAM on a large, diverse corpus of cellular imaging data, enabling it to achieve state-of-the-art (SOTA) performance across ten datasets. We also evaluated CellSAM’s zero-shot performance using a held-out dataset, LIVECell ^62^, demonstrating that it substantially outperforms existing methods for zero-shot segmentation. A deployed version of CellSAM is available at https://cellsam.deepcell.org.

**Fig. 1:**
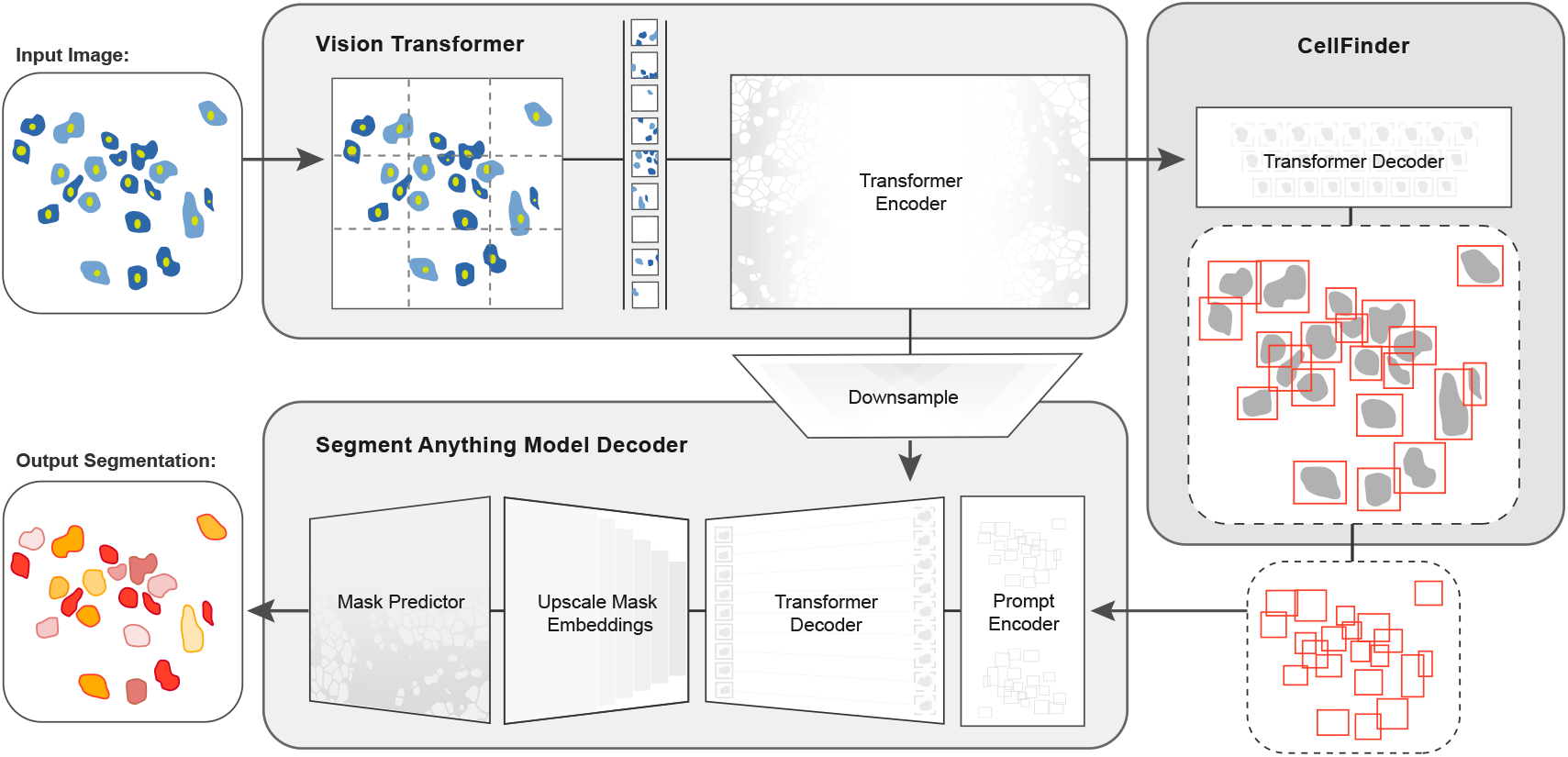
CellSAM: a foundational model for cell segmentation. CellSAM combines SAM’s mask generation and labeling capabilities with an object detection model to achieve automated inference. Input images are divided into regularly sampled patches and passed through a transformer encoder (i.e., a ViT) to generate information-rich image features. These image features are then sent to two downstream modules. The first module, CellFinder, decodes these features into bounding boxes using a transformer-based encoder-decoder pair. The second module combines these image features with prompts to generate masks using SAM’s mask decoder. CellSAM integrates these two modules using the bounding boxes generated by CellFinder as prompts for SAM. CellSAM is trained in two stages, using the pre-trained SAM model weights as a starting point. In the first stage, we train the ViT and the CellFinder model together on the object detection task. This yields an accurate CellFinder but results in a distribution shift between the ViT and SAM’s mask decoder. The second stage closes this gap by fixing the ViT and SAM mask decoder weights and fine-tuning the remainder of the SAM model (i.e., the model neck) using ground truth bounding boxes and segmentation labels.

## 2 Results

### 2.1 Construction of a dataset for general cell segmentation

A significant challenge with existing cellular segmentation methods is their inability to generalize across cellular targets, imaging modalities, and cell morphologies. To address this, we curated a dataset from the literature containing 2D images from a diverse range of targets (mammalian cells in tissues and adherent cell culture, yeast cells, bacterial cells, and mammalian cell nuclei) and imaging modalities (fluorescence, brightfield, phase contrast, and mass cytometry imaging). Our final dataset consisted of TissueNet ^28^, DeepBacs ^63^, BriFiSeg ^64^, Cellpose ^25,26^, Omnipose ^65,66^, YeastNet ^67^, YeaZ ^68^, the 2018 Kaggle Data Science Bowl dataset (DSB) ^69^, a collection of H&E datasets ^70–76^, and an internally collected dataset of phase microscopy images across eight mammalian cell lines (Phase400). We group these datasets into six types for evaluation: Tissue, Cell Culture, H&E, Bacteria, and Yeast. As the DSB ^69^ comprises cell nuclei that span several of these types, we evaluate it separately and refer to it as Nuclear, making a total of six categories for evaluation. While our method focuses on whole-cell segmentation, we included DSB ^69^ because cell nuclei are often used as a surrogate when the information necessary for whole-cell segmentation (e.g., cell membrane markers) is absent from an image. Fig. 2a shows the number of annotations per evaluation type. Finally, we used a held-out dataset LIVECell ^62^ to evaluate CellSAM’s zero-shot performance. This dataset was curated to remove low-quality images and images that did not contain sufficient information about the boundaries of closely packed cells. A detailed description of data sources and preprocessing steps can be found in Appendix A. Our full, preprocessed dataset is publicly available at https://cellsam.deepcell.org.

**Fig. 2:**
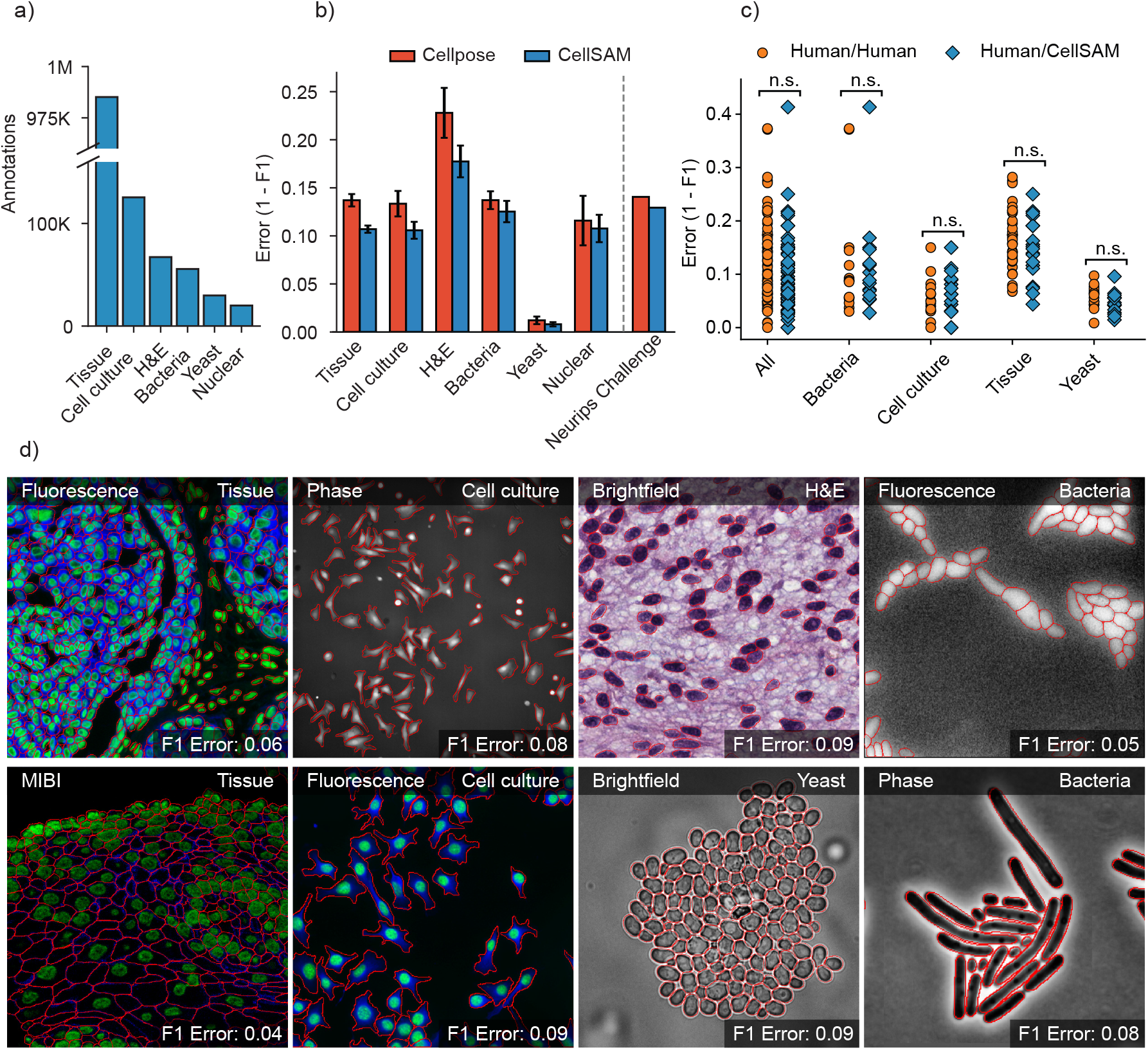
CellSAM is a strong generalist model for cell segmentation. a) For training and evaluating CellSAM, we curated a diverse cell segmentation dataset from the literature. The number of annotated cells is given for each data type. Nuclear refers to a heterogeneous dataset (DSB) ^69^ containing nuclear segmentation labels. b) Segmentation performance for CellSAM and Cellpose across different data types. We compared the segmentation error (1-F1) for models that were trained as generalists (i.e., the full dataset). Models were trained for a similar number of steps across all datasets. We observed that CellSAM-generalist had a lower error than Cellpose-generalist on all tested data categories. Furthermore, we validated this finding on a held-out competition dataset from the Weakly Supervised Cell Segmentation in Multimodality High-Resolution Microscopy Images (i.e., the NeurIPS Challenge) ^77^. Error bars were computed by computing the segmentation error per image, and then calculating the mean and standard error. c) Human vs human and CellSAM-generalist vs human (CS/human) inter-rater performance comparison. A two-sided t-test confirms that no statistical difference exists between CellSAM and human performance. d) Qualitative results of CellSAM segmentations for different data and imaging modalities. Predicted segmentations are outlined in red.

### 2.2 CellSAM creates masks using box prompts generated from CellFinder

In early experiments, we found that providing ground truth bounding boxes as prompts to SAM (ground truth prompts represent an upper bound on performance) achieved significantly higher zero-shot performance than point prompting (see Fig. S2). This is in agreement with previous analysis of SAM applied to biological ^60^ and medical ^53^ images. Because the ground-truth bounding box prompts yield accurate segmentation masks from SAM across various datasets, we sought to develop an object detector that could generate prompts for SAM in an automated fashion. Given that our zero-shot experiments demonstrated that ViT features can form robust internal representations of cellular images, we reasoned that we could build an object detector using the image features generated by SAM’s ViT. Previous work has explored this space and demonstrated that ViT backbones can achieve SOTA performance on natural images ^78,79^. For our object detection module, we use the Anchor DETR ^61^ framework with the same ViT backbone as the SAM module; we call this object detection module CellFinder. Anchor DETR is well-suited for object detection in cellular images because it formulates object detection as a set prediction task. This allows it to perform cell segmentation in images with densely packed objects, a common occurrence in cellular imaging data. Alternative bounding box methods (e.g., the R-CNN family) rely on non-maximum suppression ^80,81^, leading to poor performance in this regime. Methods that frame cell segmentation as a dense, pixel-wise prediction task (e.g., Mesmer ^28^, Cellpose ^25^, and Hover-net ^30^) assume that each pixel can be uniquely assigned to a single cell and cannot handle overlapping objects.

The ground-truth prompting scheme by itself does not achieve real-world performance standards. Our analysis showed that SAM cannot accurately segment many cell types, likely due to the distribution of images seen during training. To adapt CellSAM from natural images to cellular images, we fine-tune the SAM model neck (the layers connecting SAM’s ViT to its decoder) while leaving other layers frozen to retain generalization ability. Training CellSAM in this manner, achieved state-of-the-art accuracy when provided with ground-truth bounding box prompts.

We train CellSAM in two stages; the full details can be found in the supplement. In the first stage, we train CellFinder on the object detection task. We convert the ground truth cell masks into bounding boxes and train the ViT backbone and the CellFinder module. Once CellFinder is trained, we freeze the model weights of the ViT and fine-tune the SAM module as described above. This accounts for the distribution shifts in the ViT features that occur during the CellFinder training. Once training is complete, we use CellFinder to prompt SAM’s mask decoder. We refer to the collective method as CellSAM; Fig. 1 outlines an image’s full path through CellSAM during inference.

### 2.3 Benchmarking CellSAM’s performance on numerous biological datasets

We benchmarked CellSAM’s performance using F1 error (1-F1) as metric (Fig. 2b) against Cellpose, a widely used cell segmentation algorithm. Because our work includes both dataset and model development, we chose benchmarks that allow us to measure the contributions of data and model architecture to overall performance. Our benchmarks include comparisons to a pre-trained generalist Cellpose model (cyto3), an internally trained generalist Cellpose model, and a suite of internally trained specialist (i.e., trained on a single dataset) Cellpose models. Internally trained models were trained on the CellSAM dataset or a suitable subset using previously published training recipes, while evaluations were performed on a held-out split of the same dataset. We further evaluated CellSAM’s performance on the evaluation split of the NeurIPS Cell Segmentation challenge ^82^ (Fig. 2b). For this evaluation, we fine-tuned CellSAM with an additional hematology dataset, which was a substantial fraction of the NeurIPS challenge dataset. In almost every comparison, we found that CellSAM outperformed generalist Cellpose models (whether pre-trained or internally trained) and was equivalent to specialist Cellpose models trained exclusively on individual datasets. We highlight features of our benchmarking analyses below.

- **CellSAM is a strong generalist model.** Generalization across cell morphologies and imaging datasets has been a significant challenge for deep learning-based cell segmentation algorithms. To evaluate CellSAM’s generalization capabilities, we compared the performance of CellSAM and Cellpose models trained as specialists (i.e., on a single dataset) to generalists (i.e., on all datasets). Consistent with the literature, we observe that Cellpose’s performance degraded when trained as a generalist (Fig. S5). In contrast, we found that the performance of CellSAM-generalist was equivalent or better than CellSAM-specialist across all data categories and datasets (Fig. S4). Moreover, CellSAM-generalist out performed Cellpose-generalist in all data categories (Fig. 2b). This analysis highlights an essential feature of a foundational model: maintaining performance with increasing data diversity and scale.
- **CellSAM achieves human-level accuracy for generalized cell segmentation.** We use the error (1-F1) to assess the consistency of segmentation predictions and annotator masks across a series of images. We compared the annotations of three experts with each other (human vs. human) and with CellSAM (human vs. CellSAM). This comparison explores whether CellSAM’s performance is within the error margin created by annotator preferences (e.g., the thickness of a cell boundary). We compared annotations across four data categories: mammalian cells in tissue, mammalian cells in cell culture, bacterial cells, and yeast cells. A two-sided t-test revealed no significant differences between these two comparisons, indicating that CellSAM’s outputs are comparable to expert human annotators (Fig. 2c). This is demonstrated by non-significant p-values between CellSAM-annotator and inter-annotator agreements, specifically for Tissue: p=0.18, Cell culture: p=0.49, Yeast: p=0.11, Bacteria: p=0.90.
- **CellSAM enables fast and accurate labeling.** When provided with ground truth bounding boxes, CellSAM achieves high-quality cell masks without any fine-tuning on unseen datasets (Figure S6). Because drawing bounding boxes consumes significantly less time than drawing individual masks, this means CellSAM can be used to generate highly accurate labels quickly, even for out-of-distribution data.
- **CellSAM is a strong zero-shot and few-shot learner.** We used the LIVECell dataset to explore CellSAM’s performance in zero-shot and few-shot settings. We stratified CellSAM’s zero-shot by cell lines present in LIVECell (Supplemental Fig. S6b). We found that while performance varied by cell line, we could recover adequate performance in the few-shot regime for a number of the cell lines (e.g., *A172*). Supplemental Figures S6 and S7 show that CellSAM improves its performance with only ten additional fields of view (10^2^ − 10^3^ cells) for each cell line. We found fine-tuning could not recover performance for cell lines with morphologies far from the training data distribution (e.g., *SHSY5Y*). This may reflect a limitation of bounding boxes as a prompting strategy for SAM models.

### 2.4 CellSAM unifies biological image analysis workflows

Cell segmentation is a critical component of many spatial biology analysis pipelines; a single foundation model that generalizes across cell morphologies and imaging methods would fill a crucial gap in modern biological workflows by expanding the scope of the data that can be processed. In this section, we demonstrate how the same CellSAM-generalist model (not finetuned to any particular dataset) can be used across biological imaging pipelines by highlighting two use cases – spatial transcriptomics and live-cell imaging (Fig.3).

Spatial transcriptomics methods measure single-cell gene expression while retaining the spatial organization of the sample. These experiments (e.g., MERFISH ^83^ and seqFISH ^84^) fluorescently label individual mRNA transcripts; the number of spots for a gene inside a cell corresponds to that gene’s expression level in that cell. These methods enable the investigation of spatial gene expression patterns from the sub-cellular to tissue scales but require accurate cell segmentation to yield meaningful insights. Here, we use CellSAM-generalist in combination with Polaris ^85^, a deep learning-enabled analysis pipeline for image-based spatial transcriptomics, to analyze gene expression at the single-cell level in MERFISH ^86^ and seqFISH ^84^ data (Fig. 3). With accurate segmentation, we can assign genes to specific cells (Fig. 3). We note that CellSAM can perform segmentation on either images of nuclear and membrane stains or images derived from the spots themselves (e.g., a maximum intensity projection of all spots). CellSAM’s ability to perform nuclear and whole-cell segmentation for challenging tissue images of dense cells with complex morphologies expands the scope of datasets to which Polaris can be applied.

**Fig. 3:**
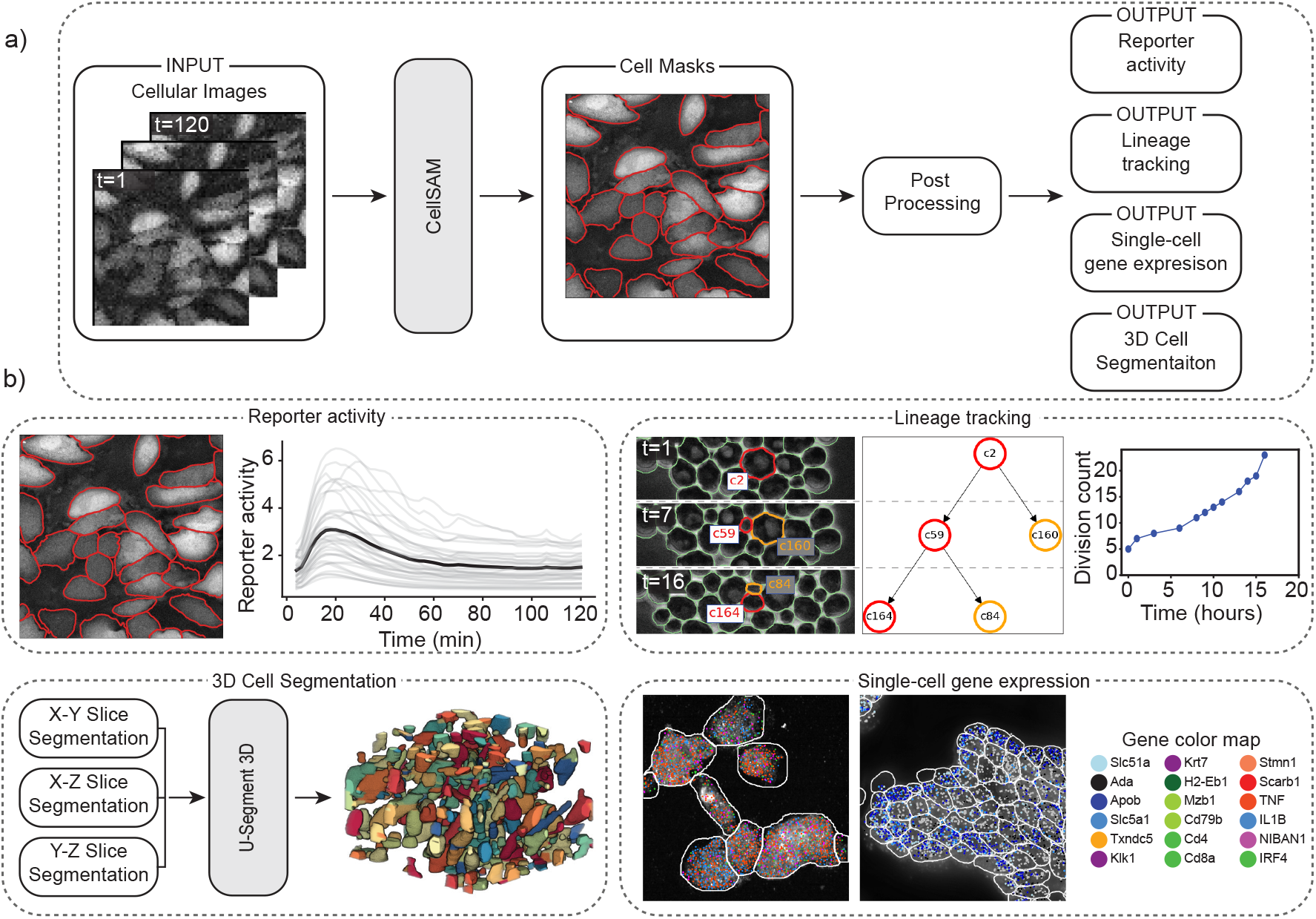
CellSAM unifies biological imaging analysis workflows. Because CellSAM-generalist functions across image modalities and cellular targets, it can be immediately applied across bioimaging analysis workflows without requiring task-specific adaptations. (a) We schematically depict how CellSAM-generalist fits into the analysis pipeline for live cell imaging and spatial transcriptomics, eliminating the need for different segmentation tools and expanding the scope of possible assays to which these tools can be applied. (b) Here, we demonstrate the application of CellSAM-generalist to various biological imaging tasks. (Top Left) Segmentations from CellSAM are used to track cells ^87^ and quantify fluorescent live-cell reporter activity in cell culture. (Top Right) CellSAM segments cells in multiple frames from a video of budding yeast cells. These cells are tracked across frames using a tracking algorithm ^87^ that ensures consistent identities, enabling accurate lineage construction and cell division quantification. (Bottom Left) CellSAM is used to segment slices of a 3D image, and these segmented slices are fed into a 3D U-Net ^89^ to create a 3D segmentation. (Bottom Right) Segmentations generated using CellSAM are integrated with Polaris ^85^, a spatial transcriptomics analysis pipeline. Because of CellSAM ‘s generalist nature, we can apply this workflow across sample types (e.g., tissue and cell culture) and imaging modalities (e.g., seqFISH and MERFISH). Datasets of cultured macrophage cells (seqFISH) and mouse ileum tissue (MERFISH) ^86^ were used to generate the data in this example. MERFISH segmentations were generated with CellSAM with an image of a nuclear and membrane stain; seqFISH segmentations were generated with CellSAM with a maximum intensity projection image of all spots.

The dynamics of cellular systems captured by live-cell imaging experiments elucidate various cellular processes such as cell division, morphological transitions, and signal transduction ^34^. The analysis of live-cell imaging data requires segmenting and tracking individual cells throughout whole movies. Here, we use CellSAM in combination with a cell tracking algorithm ^87^ (Fig. 3) in two settings. The first was a live cell imaging experiment with HeLa cells transiently expressing an AMP Kinase reporter ^88^ dosed with 20mM 2-Deoxy-D-glucose, a setup reflective of many experiments exploring cell signaling dynamics ^13^. We imaged, segmented, and tracked the cells over 60 frames or 120 minutes to quantify AMP Kinase activity over time (Fig. 3). The second setting was lineage tracking in budding yeast cells. We again used CellSAM and cell tracking to segment and track cells; we further used a division detection algorithm to count the cumulative number of divisions over time and trace individual cell lineages (Fig. 3).

CellSAM can be used to segment 3D images. We developed a workflow that generates 3D segmentations by segmenting individual slices and aggregating them to 3D using the u-Segment3D ^89^ algorithm (Fig. 3b). We demonstrate on a thin slice of epidermal organoid ^89^, as well as EASI-FISH from the lateral hypothalamus ^90^.

CellSAM’s generality ensures consistent performance; in our experiments, we did not encounter catastrophic failure modes. This is in contrast to specialist models, where failures from inference on out-of-distribution data are common. CellSAM thus enables the analysis of data modalities for which specialist models do not exist (including any modality without widespread data). Furthermore, we simplify the analysis pipeline by eliminating the need for a large collection of potentially fragile specialist models. We note that use cases are representative of the modularity of bio-image analysis pipelines; most pipelines can be broken up into a few key steps. As CellSAM demonstrates, as the algorithms that perform these steps generalize, so does the entire pipeline.

## 3 Discussion

Cell segmentation is a critical task for cellular imaging experiments. While deep learning methods have made substantial progress in recent years, there remains a need for methods that can generalize across diverse images and further reduce the marginal cost of image labeling. In this work, we sought to meet these needs by developing CellSAM, a foundation model for cell segmentation. Transformer-based methods for cell segmentation have shown promising performance^77^. CellSAM builds on these works by integrating the mask generation capabilities of SAM with transformer-based object detection to empower both scalable image labeling and automated inference. We trained CellSAM on a diverse dataset, and our benchmarking demonstrated that CellSAM achieves human-level performance on generalized cell segmentation. Compared to previous methods, CellSAM preserves its performance when trained on increasingly diverse data, which is essential for a foundational model. We found that CellSAM could be used on novel cell types in a zero-shot setting, and that re-training with few labels could yield a strong boost in performance if needed. Moreover, we demonstrated that CellSAM’s ability to generalize can be extended to entire image analysis pipelines, as illustrated by use cases in spatial transcriptomics and live cell imaging. Given its utility in image labeling and high accuracy during inference, we believe CellSAM is a valuable contribution to the field, both as a tool for spatial biology and as a means to creating the data infrastructure required for cellular imaging’s AI-powered future. To facilitate the former, we have deployed a user interface for CellSAM at https://cellsam.deepcell.org/ that allows for both automated and manual prompting. We also created a Napari plugin so that CellSAM can easily be integrated into existing workflows.

The work described here is important beyond aiding life scientists with cell segmentation. First, foundation models are immensely useful for natural language and vision tasks and hold similar promise for the life sciences, provided they are suitably adapted to this new domain. We can see several uses for CellSAM that might be within reach of future work. Much like what has occurred with natural images, we foresee that the integration of natural language labels in addition to cell-level labels might lead to vision-language models capable of generating human-like descriptors of cellular images with entity-level resolution ^48,91^. More powerful generalization capabilities may enable the standardization of cellular image analysis pipelines across all the life sciences. If the accuracy is sufficient, microbiologists and tissue biologists could use the same collection of foundation models for interpreting their imaging data even for challenging experiments^92,93^.

While the work presented here highlights the potential foundation models hold for cellular image analysis, much work remains to be done for this future to manifest. Extension of this methodology to 3D imaging data is essential; recent work on memory-efficient attention kernels ^94^ will aid these efforts. Exploring how to enable foundation models to leverage the full information content of images (e.g., multiple stains, temporal information for movies, etc.) is an essential avenue of future work. Expanding the space of labeled data remains a priority - this includes images of perturbed cells and cells with more challenging morphologies (e.g., neurons). Data generated by pooled optical screens ^95^ may synergize well with the data needs of foundation models. Compute-efficient fine-tuning strategies must be developed to enable flexible adaptation to new image domains. Lastly, prompt engineering is a critical area of future work, as it is essential to maximizing model performance. The work we presented here can be thought of as prompt engineering, as we leverage CellFinder to produce bounding box prompts for SAM. As more challenging labeled datasets are incorporated, the nature of the “best” prompts will likely evolve. Finding the best prompts for these new data is a task that will likely fall on both the computer vision and life science communities.

## Declarations

## Code Availability

The code for CellSAM is publicly available at https://github.com/vanvalenlab/cellsam.

## Data Availability

All datasets with test/training/validation splits are publicly available at https://cellsam.deepcell.org.

## Author Contributions

UI, MM contributed equally to this project and have the right to list themselves first in bibliographic documents. RD, QL contributed equally to this project and have the right to list themselves second in bibliographic documents. UI, MM, YY, and DVV conceived the project; UI, MM, QL, YY, and DVV performed algorithm design for CellFinder and CellSAM; MM implemented the CellSAM architecture; UI, MM, and QL implemented CellFinder. UI, MM, and QL carried out the experiments and evaluations of the method. GG and PP provided input for developing CellFinder; QL and UI performed model benchmarking; QL and RD developed data pipelines, RD developed the computational infrastructure for model training; RD, EP, EP, MS, QL, CY, and EL performed data engineering; EL, CY, and UI performed CellSAM integration with bioimaging workflows. MM performed 3D CellSAM integration. AA, MA, CB performed annotations on images for human-human comparison. RB and DVV supervised the software engineering, DVV supervised the project.

## Acknowledgements

We thank Leeat Keren, Noah Greenwald, Sam Cooper, Jan Funke, Uri Manor, Joe Horsman, Michael Baym, Paul Blainey, Ian Cheeseman, Manuel Leonetti, Neehar Kondapaneni, and Elijah Cole for valuable conversations and insightful feedback. We thank Felix Zhou for helping with the 3D segmentation. We also thank William Graf, Geneva Miller, and Kevin Yu, whose time in the Van Valen lab established the infrastructure and software tools that made this work possible. We thank Nader Khalil, Harper Carroll, Alec Fong, and the entire Brev.dev team for their support in establishing the computational infrastructure required for this work. We also thank Rosalind J. Xu and Jeffrey Moffitt for providing unpublished MERFISH data for the spatial transcriptomics workflow. We utilized images of the HeLa cell line in this research. Henrietta Lacks and the HeLa cell line established from her tumor cells without her knowledge or consent in 1951 has significantly contributed to scientific progress and advances in human health. We are grateful to Lacks, now deceased, and the Lacks family for their contributions to biomedical research. This work was supported by awards from the Shurl and Kay Curci Foundation (to DVV), the Rita Allen Foundation (to DVV), the Susan E. Riley Foundation (to DVV), the Pew-Stewart Cancer Scholars program (to DVV), the Gordon and Betty Moore Foundation (to DVV), the Schmidt Academy for Software Engineering (to SL), the Michael J. Fox Foundation through the Aligning Science Across Parkinson’s consortium (to DVV), the Heritage Medical Research Institute (to DVV), the National Institutes of Health New Innovator program (DP2-GM149556) (to DVV), the National Institutes of Health HuBMAP consortium (OT2-OD033756) (to DVV), and the Howard Hughes Medical Institute Freeman Hrabowski Scholars program (to DVV).

National Institutes of Health (R01-MH123612A) (to PP). NIH/Ohio State University (R01-DC014498) (to PP). Chen Institute (to PP). The Emerald Foundation and Black in Cancer (to UI). Caltech Presidential Postdoctoral Fellowship Program (PPFP) (to UI).

## References

[1] G. Palla, D. S. Fischer, A. Regev, and F. J. Theis, “Spatial components of molecular tissue biology,” Nature Biotechnology, vol. 40, no. 3, pp. 308–318, 2022.

[2] J. R. Moffitt, E. Lundberg, and H. Heyn, “The emerging landscape of spatial profiling technologies,” Nature Reviews Genetics, vol. 23, no. 12, pp. 741–759, 2022.

[3] L. Moses and L. Pachter, “Museum of spatial transcriptomics,” Nature Methods, vol. 19, no. 5, pp. 534–546, 2022.

[4] J. W. Hickey, E. K. Neumann, A. J. Radtke, J. M. Camarillo, R. T. Beuschel, A. Albanese, E. McDonough, J. Hatler, A. E. Wiblin, J. Fisher et al., “Spatial mapping of protein composition and tissue organization: a primer for multiplexed antibody-based imaging,” Nature methods, vol. 19, no. 3, pp. 284–295, 2022.

[5] J. Ko, M. Wilkovitsch, J. Oh, R. H. Kohler, E. Bolli, M. J. Pittet, C. Vinegoni, D. B. Sykes, H. Mikula, R. Weissleder et al., “Spatiotemporal multiplexed immunofluorescence imaging of living cells and tissues with bioorthogonal cycling of fluorescent probes,” Nature Biotechnology, vol. 40, no. 11, pp. 1654–1662, 2022.

[6] S. Wang, L. Furchtgott, K. C. Huang, and J. W. Shaevitz, “Helical insertion of peptidoglycan produces chiral ordering of the bacterial cell wall,” Proceedings of the National Academy of Sciences, vol. 109, no. 10, pp. E595–E604, 2012.

[7] E. R. Rojas, G. Billings, P. D. Odermatt, G. K. Auer, L. Zhu, A. Miguel, F. Chang, D. B. Weibel, J. A. Theriot, and K. C. Huang, “The outer membrane is an essential load-bearing element in gram-negative bacteria,” Nature, vol. 559, no. 7715, pp. 617–621, 2018.

[8] A. S. Hansen and E. K. O’Shea, “Limits on information transduction through amplitude and frequency regulation of transcription factor activity,” Elife, vol. 4, p. e06559, 2015.

[9] A. S. Hansen and E. K. O’shea, “Promoter decoding of transcription factor dynamics involves a trade-off between noise and control of gene expression,” Molecular systems biology, vol. 9, no. 1, p. 704, 2013.

[10] S. Tay, J. J. Hughey, T. K. Lee, T. Lipniacki, S. R. Quake, and M. W. Covert, “Single-cell nf-κb dynamics reveal digital activation and analogue information processing,” Nature, vol. 466, no. 7303, pp. 267–271, 2010.

[11] S. Regot, J. J. Hughey, B. T. Bajar, S. Carrasco, and M. W. Covert, “High-sensitivity measurements of multiple kinase activities in live single cells,” Cell, vol. 157, no. 7, pp. 1724–1734, 2014.

[12] J. Selimkhanov, B. Taylor, J. Yao, A. Pilko, J. Albeck, A. Hoffmann, L. Tsimring, and R. Wollman, “Accurate information transmission through dynamic biochemical signaling networks,” Science, vol. 346, no. 6215, pp. 1370–1373, 2014.

[13] J. E. Purvis and G. Lahav, “Encoding and decoding cellular information through signaling dynamics,” Cell, vol. 152, no. 5, pp. 945–956, 2013.

[14] M. Alieva, A. K. Wezenaar, E. J. Wehrens, and A. C. Rios, “Bridging live-cell imaging and next-generation cancer treatment,” Nature Reviews Cancer, pp. 1–15, 2023.

[15] J. Cao, G. Guan, V. W. S. Ho, M.-K. Wong, L.-Y. Chan, C. Tang, Z. Zhao, and H. Yan, “Establishment of a morphological atlas of the caenorhabditis elegans embryo using deep-learning-based 4d segmentation,” Nature communications, vol. 11, no. 1, p. 6254, 2020.

[16] D. L. Schmitt, S. Mehta, and J. Zhang, “Study of spatiotemporal regulation of kinase signaling using genetically encodable molecular tools,” Current opinion in chemical biology, vol. 71, p. 102224, 2022.

[17] K. A. Earle, G. Billings, M. Sigal, J. S. Lichtman, G. C. Hansson, J. E. Elias, M. R. Amieva, K. C. Huang, and J. L. Sonnenburg, “Quantitative imaging of gut microbiota spatial organization,” Cell host & microbe, vol. 18, no. 4, pp. 478–488, 2015.

[18] J. M. Peters, A. Colavin, H. Shi, T. L. Czarny, M. H. Larson, S. Wong, J. S. Hawkins, C. H. Lu, B.-M. Koo, E. Marta et al., “A comprehensive, crispr-based functional analysis of essential genes in bacteria,” Cell, vol. 165, no. 6, pp. 1493–1506, 2016.

[19] S. Jun, F. Si, R. Pugatch, and M. Scott, “Fundamental principles in bacterial physiology—history, recent progress, and the future with focus on cell size control: a review,” Reports on Progress in Physics, vol. 81, no. 5, p. 056601, 2018.

[20] P. Wang, L. Robert, J. Pelletier, W. L. Dang, F. Taddei, A. Wright, and S. Jun, “Robust growth of escherichia coli,” Current biology, vol. 20, no. 12, pp. 1099–1103, 2010.

[21] W.-K. Huh, J. V. Falvo, L. C. Gerke, A. S. Carroll, R. W. Howson, J. S. Weissman, and E. K. O’Shea, “Global analysis of protein localization in budding yeast,” Nature, vol. 425, no. 6959, pp. 686–691, 2003.

[22] N. Hao, B. A. Budnik, J. Gunawardena, and E. K. O’Shea, “Tunable signal processing through modular control of transcription factor translocation,” Science, vol. 339, no. 6118, pp. 460–464, 2013.

[23] N. Hao and E. K. O’shea, “Signal-dependent dynamics of transcription factor translocation controls gene expression,” Nature structural & molecular biology, vol. 19, no. 1, pp. 31–39, 2012.

[24] M. Schwartz, U. Israel, X. Wang, E. Laubscher, C. Yu, R. Dilip, Q. Li, J. Mari, J. Soro, K. Yu et al., “Scaling biological discovery at the interface of deep learning and cellular imaging,” Nature Methods, vol. 20, no. 7, pp. 956–957, 2023.

[25] C. Stringer, T. Wang, M. Michaelos, and M. Pachitariu, “Cellpose: a generalist algorithm for cellular segmentation,” Nature methods, vol. 18, no. 1, pp. 100–106, 2021.

[26] M. Pachitariu and C. Stringer, “Cellpose 2.0: how to train your own model,” Nature Methods, pp. 1–8, 2022.

[27] U. Schmidt, M. Weigert, C. Broaddus, and G. Myers, “Cell detection with star-convex polygons,” in Medical Image Computing and Computer Assisted Intervention–MICCAI 2018: 21st International Conference, Granada, Spain, September 16-20, 2018, Proceedings, Part II 11. Springer, 2018, pp. 265–273.

[28] N. F. Greenwald, G. Miller, E. Moen, A. Kong, A. Kagel, T. Dougherty, C. C. Fullaway, B. J. McIntosh, K. X. Leow, M. S. Schwartz et al., “Whole-cell segmentation of tissue images with human-level performance using large-scale data annotation and deep learning,” Nature biotechnology, vol. 40, no. 4, pp. 555–565, 2022.

[29] R. Hollandi, A. Szkalisity, T. Toth, E. Tasnadi, C. Molnar, B. Mathe, I. Grexa, J. Molnar, A. Balind, M. Gorbe et al., “nucleaizer: a parameter-free deep learning framework for nucleus segmentation using image style transfer,” Cell Systems, vol. 10, no. 5, pp. 453–458, 2020.

[30] S. Graham, Q. D. Vu, S. E. A. Raza, A. Azam, Y. W. Tsang, J. T. Kwak, and N. Rajpoot, “Hover-net: Simultaneous segmentation and classification of nuclei in multi-tissue histology images,” Medical image analysis, vol. 58, p. 101563, 2019.

[31] W. Wang, D. A. Taft, Y.-J. Chen, J. Zhang, C. T. Wallace, M. Xu, S. C. Watkins, and J. Xing, “Learn to segment single cells with deep distance estimator and deep cell detector,” Computers in biology and medicine, vol. 108, pp. 133–141, 2019.

[32] C. Stringer and M. Pachitariu, “Cellpose3: one-click image restoration for improved cellular segmentation,” bioRxiv, pp. 2024–02, 2024.

[33] D. A. Van Valen, T. Kudo, K. M. Lane, D. N. Macklin, N. T. Quach, M. M. DeFelice, I. Maayan, Y. Tanouchi, E. A. Ashley, and M. W. Covert, “Deep learning automates the quantitative analysis of individual cells in live-cell imaging experiments,” PLoS computational biology, vol. 12, no. 11, p. e1005177, 2016.

[34] M. S. Schwartz, E. Moen, G. Miller, T. Dougherty, E. Borba, R. Ding, W. Graf, E. Pao, and D. V. Valen, “Caliban: Accurate cell tracking and lineage construction in live-cell imaging experiments with deep learning,” bioRxiv, 2023. [Online]. Available: https://www.biorxiv.org/content/early/2023/09/12/803205

[35] A. Vaswani, N. Shazeer, N. Parmar, J. Uszkoreit, L. Jones, A. N. Gomez, L. Kaiser, and I. Polosukhin, “Attention is all you need,” Advances in neural information processing systems, vol. 30, 2017.

[36] R. Bommasani, D. A. Hudson, E. Adeli, R. Altman, S. Arora, S. von Arx, M. S. Bernstein, J. Bohg, A. Bosselut, E. Brunskill et al., “On the opportunities and risks of foundation models,” arXiv preprint arXiv:2108.07258, 2021.

[37] T. Brown, B. Mann, N. Ryder, M. Subbiah, J. D. Kaplan, P. Dhariwal, A. Neelakantan, P. Shyam, G. Sastry, A. Askell et al., “Language models are few-shot learners,” Advances in neural information processing systems, vol. 33, pp. 1877–1901, 2020.

[38] OpenAI, “Gpt-4 technical report,” 2023.

[39] Z. Lin, H. Akin, R. Rao, B. Hie, Z. Zhu, W. Lu, N. Smetanin, R. Verkuil, O. Kabeli, Y. Shmueli et al., “Evolutionary-scale prediction of atomic-level protein structure with a language model,” Science, vol. 379, no. 6637, pp. 1123–1130, 2023.

[40] N. Brandes, D. Ofer, Y. Peleg, N. Rappoport, and M. Linial, “Proteinbert: a universal deep-learning model of protein sequence and function,” Bioinformatics, vol. 38, no. 8, pp. 2102–2110, 2022.

[41] A. Elnaggar, M. Heinzinger, C. Dallago, G. Rehawi, Y. Wang, L. Jones, T. Gibbs, T. Feher, C. Angerer, M. Steinegger, D. Bhowmik, and B. Rost, “Prottrans: Towards cracking the language of life’s code through self-supervised learning,” bioRxiv, 2021. [Online]. Available: https://www.biorxiv.org/content/early/2021/05/04/2020.07.12.199554

[42] A. Madani, B. McCann, N. Naik, N. S. Keskar, N. Anand, R. R. Eguchi, P.-S. Huang, and R. Socher, “Progen: Language modeling for protein generation,” arXiv preprint arXiv:2004.03497, 2020.

[43] D. J. Beal, “Esm 2.0: State of the art and future potential of experience sampling methods in organizational research,” Annu. Rev. Organ. Psychol. Organ. Behav., vol. 2, no. 1, pp. 383–407, 2015.

[44] A. Dosovitskiy, L. Beyer, A. Kolesnikov, D. Weissenborn, X. Zhai, T. Unterthiner, M. Dehghani, M. Minderer, G. Heigold, S. Gelly et al., “An image is worth 16×16 words: Transformers for image recognition at scale,” arXiv preprint arXiv:2010.11929, 2020.

[45] M. Caron, H. Touvron, I. Misra, H. Jégou, J. Mairal, P. Bojanowski, and A. Joulin, “Emerging properties in self-supervised vision transformers,” in Proceedings of the IEEE/CVF international conference on computer vision, 2021, pp. 9650–9660.

[46] M. Oquab, T. Darcet, T. Moutakanni, H. Vo, M. Szafraniec, V. Khalidov, P. Fernandez, D. Haziza, F. Massa, A. El-Nouby et al., “Dinov2: Learning robust visual features without supervision,” arXiv preprint arXiv:2304.07193, 2023.

[47] Y. Fang, W. Wang, B. Xie, Q. Sun, L. Wu, X. Wang, T. Huang, X. Wang, and Y. Cao, “Eva: Exploring the limits of masked visual representation learning at scale,” in Proceedings of the IEEE/CVF Conference on Computer Vision and Pattern Recognition, 2023, pp. 19358–19369.

[48] A. Radford, J. W. Kim, C. Hallacy, A. Ramesh, G. Goh, S. Agarwal, G. Sastry, A. Askell, P. Mishkin, J. Clark et al., “Learning transferable visual models from natural language supervision,” in International conference on machine learning. PMLR, 2021, pp. 8748–8763.

[49] J.-B. Alayrac, J. Donahue, P. Luc, A. Miech, I. Barr, Y. Hasson, K. Lenc, A. Mensch, K. Millican, M. Reynolds et al., “Flamingo: a visual language model for few-shot learning,” Advances in Neural Information Processing Systems, vol. 35, pp. 23716–23736, 2022.

[50] D. Hernandez, J. Kaplan, T. Henighan, and S. McCandlish, “Scaling laws for transfer,” arXiv preprint arXiv:2102.01293, 2021.

[51] X. Zhai, A. Kolesnikov, N. Houlsby, and L. Beyer, “Scaling vision transformers,” in Proceedings of the IEEE/CVF conference on computer vision and pattern recognition, 2022, pp. 12104–12113.

[52] A. Kirillov, E. Mintun, N. Ravi, H. Mao, C. Rolland, L. Gustafson, T. Xiao, S. Whitehead, A. C. Berg, W.-Y. Lo et al., “Segment anything,” arXiv preprint arXiv:2304.02643, 2023.

[53] Y. Huang, X. Yang, L. Liu, H. Zhou, A. Chang, X. Zhou, R. Chen, J. Yu, J. Chen, C. Chen et al., “Segment anything model for medical images?” Medical Image Analysis, vol. 92, p. 103061, 2024.

[54] Y. Zhang, T. Zhou, S. Wang, P. Liang, and D. Z. Chen, “Input augmentation with sam: Boosting medical image segmentation with segmentation foundation model,” 2023.

[55] W. Lei, X. Wei, X. Zhang, K. Li, and S. Zhang, “Medlsam: Localize and segment anything model for 3d medical images,” 2023.

[56] P. Shi, J. Qiu, S. M. D. Abaxi, H. Wei, F. P.-W. Lo, and W. Yuan, “Generalist vision foundation models for medical imaging: A case study of segment anything model on zero-shot medical segmentation,” Diagnostics, vol. 13, no. 11, p. 1947, 2023.

[57] M. Hu, Y. Li, and X. Yang, “Skinsam: Empowering skin cancer segmentation with segment anything model,” 2023.

[58] R. Deng, C. Cui, Q. Liu, T. Yao, L. W. Remedios, S. Bao, B. A. Landman, L. E. Wheless, L. A. Coburn, K. T. Wilson, Y. Wang, S. Zhao, A. B. Fogo, H. Yang, Y. Tang, and Y. Huo, “Segment anything model (sam) for digital pathology: Assess zero-shot segmentation on whole slide imaging,” 2023.

[59] F. Hörst, M. Rempe, L. Heine, C. Seibold, J. Keyl, G. Baldini, S. Ugurel, J. Siveke, B. Grünwald, J. Egger, and J. Kleesiek, “Cellvit: Vision transformers for precise cell segmentation and classification,” 2023.

[60] A. Archit, S. Nair, N. Khalid, P. Hilt, V. Rajashekar, M. Freitag, S. Gupta, A. Dengel, S. Ahmed, and C. Pape, “Segment anything for microscopy,” bioRxiv, 2023. [Online]. Available: https://www.biorxiv.org/content/early/2023/08/22/2023.08.21.554208

[61] Y. Wang, X. Zhang, T. Yang, and J. Sun, “Anchor detr: Query design for transformer-based detector,” in Proceedings of the AAAI conference on artificial intelligence, vol. 36, no. 3, 2022, pp. 2567–2575.

[62] C. Edlund, T. R. Jackson, N. Khalid, N. Bevan, T. Dale, A. Dengel, S. Ahmed, J. Trygg, and R. Sjögren, “Livecell—a large-scale dataset for label-free live cell segmentation,” Nature methods, vol. 18, no. 9, pp. 1038–1045, 2021.

[63] C. Spahn, E. Gomez-de Mariscal, R. F. Laine, P. M. Pereira, L. von Chamier, M. Conduit, M. G. Pinho, G. Jacquemet, S. Holden, M. Heilemann et al., “Deepbacs for multi-task bacterial image analysis using open-source deep learning approaches,” Communications Biology, vol. 5, no. 1, p. 688, 2022.

[64] G. Mathieu, E. D. Bachir et al., “Brifiseg: a deep learning-based method for semantic and instance segmentation of nuclei in brightfield images,” arXiv preprint arXiv:2211.03072, 2022.

[65] K. J. Cutler, C. Stringer, P. A. Wiggins, and J. D. Mougous, “Omnipose: a high-precision morphology-independent solution for bacterial cell segmentation,” bioRxiv, 2021.

[66] K. J. Cutler, C. Stringer, T. W. Lo, L. Rappez, N. Stroustrup, S. Brook Peterson, P. A. Wiggins, and J. D. Mougous, “Omnipose: a high-precision morphology-independent solution for bacterial cell segmentation,” Nature methods, vol. 19, no. 11, pp. 1438–1448, 2022.

[67] H. Kim, J. Shin, E. Kim, H. Kim, S. Hwang, J. E. Shim, and I. Lee, “Yeastnet v3: a public database of data-specific and integrated functional gene networks for saccharomyces cerevisiae,” Nucleic acids research, vol. 42, o. D1, pp. D731–D736, 2014.

[68] N. Dietler, M. Minder, V. Gligorovski, A. M. Economou, D. A. H. Lucien Joly, A. Sadeghi, C. H. Michael Chan, M. Koziński, M. Weigert, A.-F. Bitbol et al., “Yeaz: A convolutional neural network for highly accurate, label-free segmentation of yeast microscopy images,” bioRxiv, pp. 2020–05, 2020.

[69] J. C. Caicedo, A. Goodman, K. W. Karhohs, B. A. Cimini, J. Ackerman, M. Haghighi, C. Heng, T. Becker, M. Doan, C. McQuin et al., “Nucleus segmentation across imaging experiments: the 2018 data science bowl,” Nature methods, vol. 16, no. 12, pp. 1247–1253, 2019.

[70] N. Kumar, R. Verma, S. Sharma, S. Bhargava, A. Vahadane, and A. Sethi, “A dataset and a technique for generalized nuclear segmentation for computational pathology,” IEEE transactions on medical imaging, vol. 36, no. 7, pp. 1550–1560, 2017.

[71] A. Mahbod, G. Schaefer, B. Bancher, C. Low, G. Dorffner, R. Ecker, and I. Ellinger, “Cryonuseg: A dataset for nuclei instance segmentation of cryosectioned h&e-stained histological images,” Computers in biology and medicine, vol. 132, p. 104349, 2021.

[72] A. Mahbod, C. Polak, K. Feldmann, R. Khan, K. Gelles, G. Dorffner, R. Woitek, S. Hatamikia, and I. Ellinger, “Nuinsseg: a fully annotated dataset for nuclei instance segmentation in h&e-stained histological images,” arXiv preprint arXiv:2308.01760, 2023.

[73] P. Naylor, M. Laé, F. Reyal, and T. Walter, “Segmentation of nuclei in histopathology images by deep regression of the distance map,” IEEE transactions on medical imaging, vol. 38, no. 2, pp. 448–459, 2018.

[74] N. Kumar, R. Verma, D. Anand, Y. Zhou, O. F. Onder, E. Tsougenis, H. Chen, P.-A. Heng, J. Li, Z. Hu et al., “A multi-organ nucleus segmentation challenge,” IEEE transactions on medical imaging, vol. 39, no. 5, pp. 1380–1391, 2019.

[75] Q. D. Vu, S. Graham, T. Kurc, M. N. N. To, M. Shaban, T. Qaiser, N. A. Koohbanani, S. A. Khurram, J. Kalpathy-Cramer, T. Zhao et al., “Methods for segmentation and classification of digital microscopy tissue images,” Frontiers in bioengineering and biotechnology, p. 53, 2019.

[76] R. Verma, N. Kumar, A. Patil, N. C. Kurian, S. Rane, S. Graham, Q. D. Vu, M. Zwager, S. E. A. Raza, N. Rajpoot et al., “Monusac2020: A multi-organ nuclei segmentation and classification challenge,” IEEE Transactions on Medical Imaging, vol. 40, no. 12, pp. 3413–3423, 2021.

[77] J. Ma, R. Xie, S. Ayyadhury, C. Ge, A. Gupta, R. Gupta, S. Gu, Y. Zhang, G. Lee, J. Kim, W. Lou, H. Li, E. Upschulte, T. Dickscheid, J. G. de Almeida, Y. Wang, L. Han, X. Yang, M. Labagnara, V. Gligorovski, M. Scheder, S. J. Rahi, C. Kempster, A. Pollitt, L. Espinosa, T. Mignot, J. M. Middeke, J.-N. Eckardt, W. Li, Z. Li, X. Cai, B. Bai, N. F. Greenwald, D. V. Valen, E. Weisbart, B. A. Cimini, T. Cheung, O. Brück, G. D. Bader, and B. Wang, “The multi-modality cell segmentation challenge: Towards universal solutions,” Nature Methods, vol. 21, p. 1103–1113, 2024.

[78] Y. Li, H. Mao, R. Girshick, and K. He, “Exploring plain vision transformer backbones for object detection,” in European Conference on Computer Vision. Springer, 2022, pp. 280–296.

[79] T.-Y. Lin, M. Maire, S. Belongie, J. Hays, P. Perona, D. Ramanan, P. Dollár, and C. L. Zitnick, “Microsoft coco: Common objects in context,” in Computer Vision–ECCV 2014: 13th European Conference, Zurich, Switzerland, September 6-12, 2014, Proceedings, Part V 13. Springer, 2014, pp. 740–755.

[80] R. Girshick, J. Donahue, T. Darrell, and J. Malik, “Rich feature hierarchies for accurate object detection and semantic segmentation,” in Proceedings of the IEEE conference on computer vision and pattern recognition, 2014, pp. 580–587.

[81] S. Ren, K. He, R. Girshick, and J. Sun, “Faster r-cnn: Towards real-time object detection with region proposal networks,” 2016.

[82] J. Ma, R. Xie, S. Ayyadhury, C. Ge, A. Gupta, R. Gupta, S. Gu, Y. Zhang, G. Lee, J. Kim et al., “The multimodality cell segmentation challenge: toward universal solutions,” Nature methods, pp. 1–11, 2024.

[83] K. H. Chen, A. N. Boettiger, J. R. Moffitt, S. Wang, and X. Zhuang, “Spatially resolved, highly multiplexed rna profiling in single cells,” Science, vol. 348, no. 6233, p. aaa6090, 2015.

[84] C.-H. L. Eng, M. Lawson, Q. Zhu, R. Dries, N. Koulena, Y. Takei, J. Yun, C. Cronin, C. Karp, G.-C. Yuan et al., “Transcriptome-scale super-resolved imaging in tissues by rna seqfish+,” Nature, vol. 568, no. 7751, pp. 235–239, 2019.

[85] E. Laubscher, X. J. Wang, N. Razin, T. Dougherty, R. J. Xu, L. Ombelets, E. Pao, W. Graf, J. R. Moffitt, Y. Yue et al., “Accurate single-molecule spot detection for image-based spatial transcriptomics with weakly supervised deep learning,” bioRxiv, 2023.

[86] V. Petukhov, R. J. Xu, R. A. Soldatov, P. Cadinu, K. Khodosevich, J. R. Moffitt, and P. V. Kharchenko, “Cell segmentation in imaging-based spatial transcriptomics,” Nature biotechnology, vol. 40, no. 3, pp. 345–354, 2022.

[87] E. Bochinski, V. Eiselein, and T. Sikora, “High-speed tracking-by-detection without using image information,” in 2017 14th IEEE international conference on advanced video and signal based surveillance (AVSS). IEEE, 2017, pp. 1–6.

[88] D. L. Schmitt, S. D. Curtis, A. C. Lyons, J.-f. Zhang, M. Chen, C. Y. He, S. Mehta, R. J. Shaw, and J. Zhang, “Spatial regulation of ampk signaling revealed by a sensitive kinase activity reporter,” Nature communications, vol. 13, no. 1, p. 3856, 2022.

[89] F. Y. Zhou, C. Yapp, Z. Shang, S. Daetwyler, Z. Marin, M. T. Islam, B. Nanes, E. Jenkins, G. M. Gihana, B.-J. Chang et al., “A general algorithm for consensus 3d cell segmentation from 2d segmented stacks,” bioRxiv, 2024.

[90] Y. Wang, M. Eddison, G. Fleishman, M. Weigert, S. Xu, T. Wang, K. Rokicki, C. Goina, F. E. Henry, A. L. Lemire et al., “Easi-fish for thick tissue defines lateral hypothalamus spatio-molecular organization,” Cell, vol. 184, no. 26, pp. 6361–6377, 2021.

[91] X. Wang, R. Dilip, Y. Bussi, C. Brown, E. Pradhan, Y. Jain, K. Yu, S. Li, M. Abt, K. Börner et al., “Generalized cell phenotyping for spatial proteomics with language-informed vision models,” bioRxiv, pp. 2024–11, 2024.

[92] S. Shah, E. Lubeck, W. Zhou, and L. Cai, “seqfish accurately detects transcripts in single cells and reveals robust spatial organization in the hippocampus,” Neuron, vol. 94, no. 4, pp. 752–758, 2017.

[93] D. Dar, N. Dar, L. Cai, and D. K. Newman, “Spatial transcriptomics of planktonic and sessile bacterial populations at single-cell resolution,” Science, vol. 373, no. 6556, p. eabi4882, 2021.

[94] E. Nguyen, M. Poli, M. Faizi, A. Thomas, C. Birch-Sykes, M. Wornow, A. Patel, C. Rabideau, S. Massaroli, Y. Bengio, S. Ermon, S. A. Baccus, and C. Ré, “Hyenadna: Long-range genomic sequence modeling at single nucleotide resolution,” 2023.

[95] D. Feldman, A. Singh, J. L. Schmid-Burgk, R. J. Carlson, A. Mezger, A. J. Garrity, F. Zhang, and P. C. Blainey, “Optical pooled screens in human cells,” Cell, vol. 179, no. 3, pp. 787–799, 2019.

[96] S. M. Pizer, E. P. Amburn, J. D. Austin, R. Cromartie, A. Geselowitz, T. Greer, B. ter Haar Romeny, J. B. Zimmerman, and K. Zuiderveld, “Adaptive histogram equalization and its variations,” Computer vision, graphics, and image processing, vol. 39, no. 3, pp. 355–368, 1987.

[97] J. Hosang, R. Benenson, and B. Schiele, “Learning non-maximum suppression,” in Proceedings of the IEEE conference on computer vision and pattern recognition, 2017, pp. 4507–4515.

[98] K. He, X. Zhang, S. Ren, and J. Sun, “Deep residual learning for image recognition,” in Proceedings of the IEEE conference on computer vision and pattern recognition, 2016, pp. 770–778.

[99] N. Carion, F. Massa, G. Synnaeve, N. Usunier, A. Kirillov, and S. Zagoruyko, “End-to-end object detection with transformers,” in Computer Vision–ECCV 2020: 16th European Conference, Glasgow, UK, August 23–28, 2020, Proceedings, Part I 16. Springer, 2020, pp. 213–229.

[100] I. Loshchilov and F. Hutter, “Decoupled weight decay regularization,” arXiv preprint arXiv:1711.05101, 2017.

[101] A. Paszke, S. Gross, F. Massa, A. Lerer, J. Bradbury, G. Chanan, T. Killeen, Z. Lin, N. Gimelshein, L. Antiga, A. Desmaison, A. Kopf, E. Yang, Z. DeVito, M. Raison, A. Tejani, S. Chilamkurthy, B. Steiner, L. Fang, J. Bai, and S. Chintala, “Pytorch: An imperative style, high-performance deep learning library,” in Advances in Neural Information Processing Systems 32. Curran Associates, Inc., 2019, pp. 8024–8035. [Online]. Available: http://papers.neurips.cc/paper/9015-pytorch-an-imperative-style-high-performance-deep-learning-library.pdf

[102] W. Falcon and The PyTorch Lightning team, “PyTorch Lightning,” Mar. 2019. [Online]. Available: https://github.com/Lightning-AI/lightning

