## Supplement for "CellSAM: A Foundation Model for Cell Segmentation"

### A Dataset Construction

To train CellSAM, we combined ten separate datasets spanning a variety of modalities: TissueNet[28], DeepBacs[63], BriFiSeg[64], Cellpose[25,26], Omnipose[65,66], YeastNet[67], YeaZ[68], the 2018 Kaggle Data Science Bowl (DSB)[69], a collection of H&E datasets[70–76], and an internally collected dataset of phase microscopy images across eight mammalian cell lines (Phase400). The LIVECell[62] dataset was held out for zero-shot/few-shot tests. Our collective dataset included images across multiple imaging modalities (brightfield, phase contrast, fluorescence, and mass cytometry), imaging targets (histology sections, yeast, cell culture, bacteria, nuclei), length scales, and morphologies. We didn't do any preprocessing and left pixel intensities untouched. We treated nuclear and whole-cell channels as green and blue channels in an RGB image, respectively, and the red channel was always blank. We moved the green channel to blue for nuclear-only datasets (i.e., BriFiSeg and DSB) to keep the blue channel always occupied.

If available, we used pre-determined train/val/test splits for each dataset; otherwise, we introduced 80-10-10% data splits. For datasets with multiple fields of view of the same object set, we required all FOVs to belong to the same split. We deferred all duplicated samples to the train split for published datasets with a pre-existing data leak (detected by pixel-wise hashing). Our assembled dataset kept all images in their original size. We followed a widely used annotation scheme for labeling our masks, with zero representing the background and unique positive integers representing different objects. While this format precludes accurate segmentation of overlapping objects, labels of this kind were not present in the dataset we compiled. We filtered out invalid cell labels if the label contained disjoint regions or if the label had only a 1-pixel height or width. The processed images with filtered annotations were used for training, validation, and testing. We conducted some additional processing for LIVECell[62]. We converted annotations from the COCO format to the same labeling format we used on the other datasets for consistency. We used Cellpose's[26] pre-processing function `livecell_ann_to_masks()` to remove overlapping regions. In addition, we noticed inconsistencies in ground truth labels as previously observed by the Cellpose team (see Fig 1.c in[26]). We thus manually inspected the LIVECell test split to divide the annotation quality into three classes - good, medium, and poor. We randomly selected images in the good split of the test set for the CellSAM few-shot learning task.

We modified our preparation pipeline for the NeurIPS challenge dataset. In addition to our standard preprocessing, we reduced bright spots by linearly re-scaling the raw pixel intensities such that the 99.9 percentiles corresponded to 1.0. We then normalized each image with Contrast Limited Adaptive Histogram Equalization (CLAHE)[96] with a kernel size of 128 pixels. Our assembled NeurIPS dataset used a fixed image size of 512 by 512 pixels. Images shorter than 512 pixels on either axis were zero-padded up to 512. For images with more than 512 pixels on either axis, we tiled them to 512 by 512 pixels with a 25% overlap and filled the empty regions with zeros. Any cropped images without valid annotations were removed. The NeurIPS training dataset includes all train/val/test splits from our standard datasets, plus NeurIPS training and tuning datasets. We used NeurIPS open test set for validation and hidden test set for performance report.

Statistics on our full dataset are in Table 1. To aid reproducibility, the dataset labels in Table 1 correspond to the labels present in our compiled dataset available at https://cellsam.deepcell.org.

| Dataset | Test | | Train | | Val | |
|---|---|---|---|---|---|---|
| | Images | Objects | Images | Objects | Images | Objects |
| nuc_seg_dsb | 56 | 2292 | 449 | 20036 | 57 | 2558 |
| YeaZ | 17 | 1917 | 303 | 22526 | 8 | 852 |
| Gendarme_BriFi | 22 | 796 | 179 | 6458 | 23 | 815 |
| YeastNet | 15 | 468 | 120 | 5988 | 15 | 894 |
| bact_fluor | 23 | 5388 | 129 | 16999 | 6 | 691 |
| bact_phase | 145 | 18400 | 224 | 25567 | 22 | 1682 |
| cellpose | 68 | 7201 | 486 | 58701 | 54 | 5978 |
| 2b_brightfield_dataset | 5 | 470 | 2 | 394 | 3 | 425 |
| 2b_fluorescence_dataset | 5 | 470 | 2 | 352 | 3 | 425 |
| 2c_e_coli | 15 | 1142 | 4 | 335 | 4 | 252 |
| 2d_1_SplineDist_dataset | 10 | 649 | 22 | 385 | 10 | 138 |
| 2d_2_b_subtilis | 10 | 649 | 70 | 3843 | 2 | 104 |
| 2e_e_coli | 12 | 688 | 21 | 1083 | 4 | 269 |
| s2_stardist | 35 | 2731 | 120 | 6656 | 8 | 476 |
| monuseg | 7 | 3822 | 37 | 24103 | 7 | 2861 |
| nuinsseg | 1 | 24 | 9 | 325 | 2 | 51 |
| tnbc | 5 | 391 | 40 | 3257 | 5 | 408 |
| cpm15 | 1 | 255 | 12 | 2048 | 2 | 602 |
| cpm17 | 16 | 1755 | 32 | 3766 | 16 | 2049 |
| kumar | 0 | 0 | 16 | 9905 | 0 | 0 |
| monusac | 21 | 3864 | 167 | 23852 | 21 | 3482 |
| tissuenet_wholecell | 330 | 132568 | 2576 | 985134 | 324 | 119915 |
| RAW264_ep_microscopy | 13 | 829 | 100 | 6968 | 13 | 1214 |
| 3T3_ep_microscopy | 4 | 658 | 32 | 4106 | 5 | 527 |
| HeLa_ep_microscopy | 4 | 494 | 32 | 2789 | 4 | 353 |
| HEK293_ep_microscopy | 2 | 583 | 17 | 4051 | 3 | 659 |
| CHO_ep_microscopy | 4 | 1002 | 29 | 6220 | 4 | 883 |
| A549_ep_microscopy | 7 | 623 | 60 | 4950 | 8 | 483 |
| PC3_ep_microscopy | 7 | 696 | 54 | 4649 | 7 | 571 |
| HeLa-S3_ep_microscopy | 13 | 3034 | 103 | 25421 | 13 | 3368 |
| neurips_fixed | 400 | Not publicly available | 1100 | 309917 | 50 | 6041 |

**Table 1**: Number of images and objects in each dataset split by test, train, and val.

### B CellSAM Architecture.

We adapted Anchor DETR[61] for the object detector for CellSAM (CellFinder). This choice was motivated by Anchor DETR being non-maximum suppression (NMS)[97] free. NMS suppresses bounding boxes with a high amount of overlap to remove duplicate detections. While this works well for natural images, cellular images often have tightly clustered objects, and NMS-based methods such as the R-CNN family[80,81] can suffer from a low recall in this setting. We replaced the Anchor DETR's ResNet[98] backbone with the vision transformer (ViT)[44] from the SAM model[52]; specifically, we used the base-sized ViT (ViT-B).

As the maximum number of cells per image is generally no more than 1000, we increased the number of queries $q$ to 3500, 3.5 times the maximum number of cells, based on Fig. 12 in DETR[99], which provided an estimate of the number of queries needed for a DETR method to detect all objects. We used one pattern $p$ for the Anchor generation as most objects in cellular detection are usually of similar scale.

*Training CellFinder*: We used a base learning rate of $10^{-4}$ for the Anchor DETR head and $10^{-5}$ for the SAM-ViT backbone. We used weight decay of $10^{-4}$ and clip norm of 0.1. We used AdamW[100] with a step-wise learning rate scheduler that drops the learning rate by a factor of 10 after the 1960th epoch. We trained CellFinder for 2800 epochs with a batch size of 4 across 8 H100 GPUs.

*Finetuning CellSAM*: After we trained CellFinder with the SAM-ViT backbone, the SAM-ViT output features were no longer aligned with the rest of the model (i.e., the prompt encoder and mask decoder). To close this distribution gap, we froze the SAM-ViT (such that it continues to function well with CellFinder) and trained the neck of the SAM model. We trained this neck using ground-truth bounding boxes as inputs and segmentation masks of individual cells as targets. We used a learning rate of $10^{-4}$ and weight decay of $10^{-4}$ for this training. We also used AdamW[100] for this training and did not clip the gradient.

#### B.1 Inference

At inference, we followed the following workflow. First, the input was passed through the Anchor DETR fine-tuned ViT-B. This resulted in an embedding dimension of 768. This embedding was then passed as an input to two parts of CellSAM, 1) the trained Anchor DETR module (CellFinder) and 2) the fine-tuned neck, which is a 2D convolutional network reducing the embedding dimensionality further to 256. The bounding box outputs of CellFinder were then sent into the prompt encoder, resulting in the prompt embedding. The prompt embeddings and neck embedding were then passed to the mask decoder, which outputs pixel-wise probabilities for the cell and another IoU-based confidence value for the prediction as a whole. This results in a tensor of shape $N \times W \times H$, where $N$ corresponds to the number of cells predicted. This tensor was processed with a sigmoid and a threshold operation, resulting in binarized images. Depending on the metric used, we either used this tensor directly with the $N$ scores (specifically for computation of the COCO AP @ 0.5 IoU), or we computed the argmax over the cell dimension $N$ to generate a tensor $W \times H$, where each pixel corresponded to a unique integer label for each cell.

*Thresholding*: Given CellSAM's model architecture, we had three different thresholds at inference time. First, we had a threshold on the bounding boxes generated by CellFinder, which we set to 0.4 across all datasets. We dynamically adjust this threshold by clustering the confidence values. We perform k-means clustering ($k = 2$) the box-confidences of all cells for a given images. We then compute the mean of both clusters ($T_\mu$, i.e. the seperation between likely non-cells and cells. We then adjust the final bounding box threshold as $T_{box} = \frac{2}{3} * 0.4 + \frac{1}{3} * T_\mu$. The resulting boxes were passed through the Mask Decoder, we had an overall mask score outputted by the IoU prediction head of the Mask Decoder, which we set to 0.5. Lastly, we thresholded the mask decoder output after applying the sigmoid function to each pixel, which we set at 0.5.

*CellSAM Postprocessing*: We used the same postprocessing steps that are used by Cellpose[32]. This consisted of hole filling and island removal for each predicted cell.

*Inference time*: Here we show the inference time in Supplemental Figures S1 as a function of the number of cells.

##### B.1.1 Model Implementation and Training

CellSAM was implemented in Pytorch[101]. For CellFinder we modified the official Anchor DETR repo[1]. For CellSAM, we modified the official Segment Anything repo[2]. We used Pytorch Lightning[102] to scale the training. Prototyping was done using NVIDIA's RTX 4090. We used machines with NVIDIA A6000s, A100s (40GB and 80GB versions), or H100s for the experiments in the paper. The overall training time was 6d 12h for training Cellfinder and 1d for subsequent fine-tuning (note fine-tuning was on 1 GPU).

[1]https://github.com/megvii-research/AnchorDETR
[2]https://github.com/facebookresearch/segment-anything

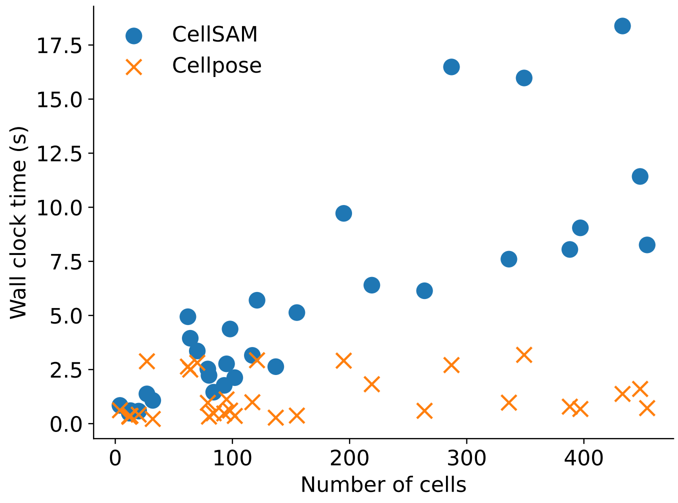


**Fig. S1**: Inference time for the CellSAM and Cellpose.

##### B.2 3D integration

We use u-segment3D[89] to generate 3D cellular segmentations. We first segment each 2D slice using the CellSAM-generalist model. We then remove small masks and fill holes using Cellpose's postprocessing. We then use the u-segment3D[89] indirect method to fuse the 2D segmentations to 3D.

#### C Benchmarking

We benchmarked the performance of CellSAM models against Cellpose[25,26] trained on our compiled datasets.

##### C.1 Cellpose Model Training.

We utilized the command provided by the Cellpose developers to train our custom Cellpose models. Using Cellpose library version 3.0.11, we trained both specialist and generalist models from scratch[26].

```
python -m cellpose --train --train_size --use_gpu --dir {} --test_dir {}
--img_filter _img --mask_filter _masks --pretrained_model None --chan 3 -chan2 2
```

We kept all the hyper-parameters untouched. We used the SGD optimizer with a weight decay of $10^{-5}$ and a batch size of 8. We trained each model for 500 epochs with a base learning rate of 0.2. We used the default learning rate scheduler. The learning rate increased linearly from 0 to 0.2 over the first ten epochs, then decreased by a factor of 2 every 10 epochs after the 400th epoch. We trained each model on a single NVIDIA A6000 GPU. In total, we trained ten specialist models and one generalist model. When evaluating the performance of the Cellpose model, we followed the implementation of the cyto3 model evaluating pipeline. We first estimated the cell diameters per image using a trained size model. Then, we predicted the segmentation masks.

##### C.2 Metrics

We used the Metrics package in the Cellpose library[25,26]. Predictions that match the ground truth labels (determined by a mask IoU $\geq 0.5$) are true positives (TP), predictions with no matching ground truth labels are false positives (FP), and ground truth labels without a valid match are false negatives (FN). For human-to-human comparisons, we used a lower threshold of 0.3 to reflect the different styles of different annotators. We computed the recall, precision, and F1 scores using the following formulas:

- Recall: recall $= \frac{\text{TP}}{\text{TP+FN}}$.
- Precision: precision $= \frac{\text{TP}}{\text{TP+FP}}$.
- F1: $F_1 = \frac{2\times\text{precision}\times\text{recall}}{\text{precision}+\text{recall}}$.

We also used the COCO evaluation metrics[79] during CellFinder's development. The COCO metrics are a widely used benchmark for assessing the object-level quality of object detection and instance segmentation methods. These metrics report Average Precision (AP), the area under the Precision-Recall curve for a given object class. In our case, we only had a single object class: cells. The AP is computed for different IoU thresholds, ranging from 0.5 to 0.95, with a step size of 0.05. We report the mean AP across all IoU thresholds, denoted as `mAP`, as well as the AP at IoU=0.5, denoted as `AP50`, to quantify CellFinder's performance. Because the object density is much higher in cellular images than in natural images, we modified the limit for the maximum number of detections from 100 to 10,000. We also fed the actual confidence score per binary prediction of the CellSAM model to the COCO evaluator. For the Cellpose models, we used a fixed confidence score of 1.0.

### D Hyperparameters

For training, there are two sets of parameters. The first is for training CellFinder , which we trained for 2800 epochs with a batch size of 4 (primarily constrained by GPU memory) across 8 GPUs. We used the AdamW optimizer with a learning rate of $10^{-5}$ for the backbone and $10^{-4}$ for the remainder of the model. We set weight decay $10^{-4}$ and dropout to 0.1. After 1960 epochs, we reduced the learning rate by a factor of 10. The full configuration file can be found in the Github repository.

The second set of parameters is fine-tuning CellSAM which we trained for 50 epochs with a cosine LR schedule. We used 3500 query positions and used the default configuration available in the GitHub repository for the rest of the parameters.

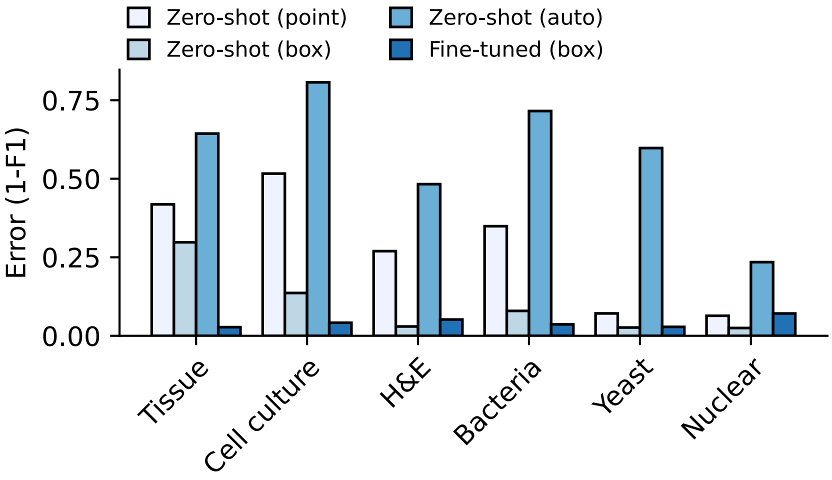


**Fig. S2**: **Preliminary prompting analysis** comparing point prompts, bounding box prompts, and SAM's auto prompting (a uniform grid of points). We used the ground truth cellular masks to generate both the point and bounding box prompts (serving as a theoretical upper bound on performance). We used the F1 error (1-F1 score) for the metric of comparison. Additionally, we used SAM with no additional training to assess zero-shot performance and compared that to a fine-tuned SAM. Here, we see the best performance was achieved using bounding box prompts with a fine-tuned model.

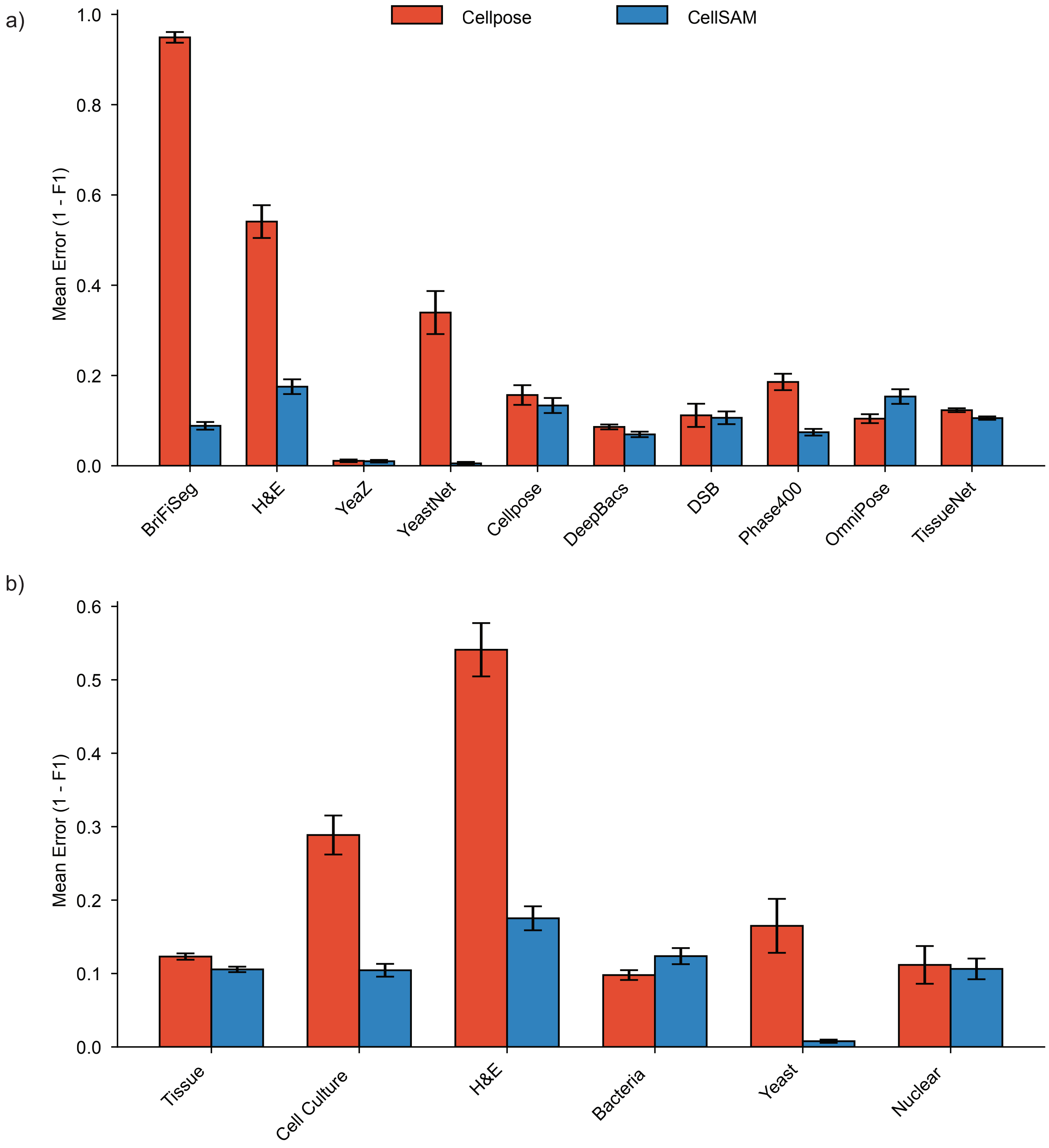


**Fig. S3**: **CellSAM-generalist Comapred to Cellpose generalist model (cyto3)**. Here, we compare CellSAM-generalist performance to Cellpose's cyto3 model. a) examines the per dataset performance of these two models. b) examples the performance across the various data types.

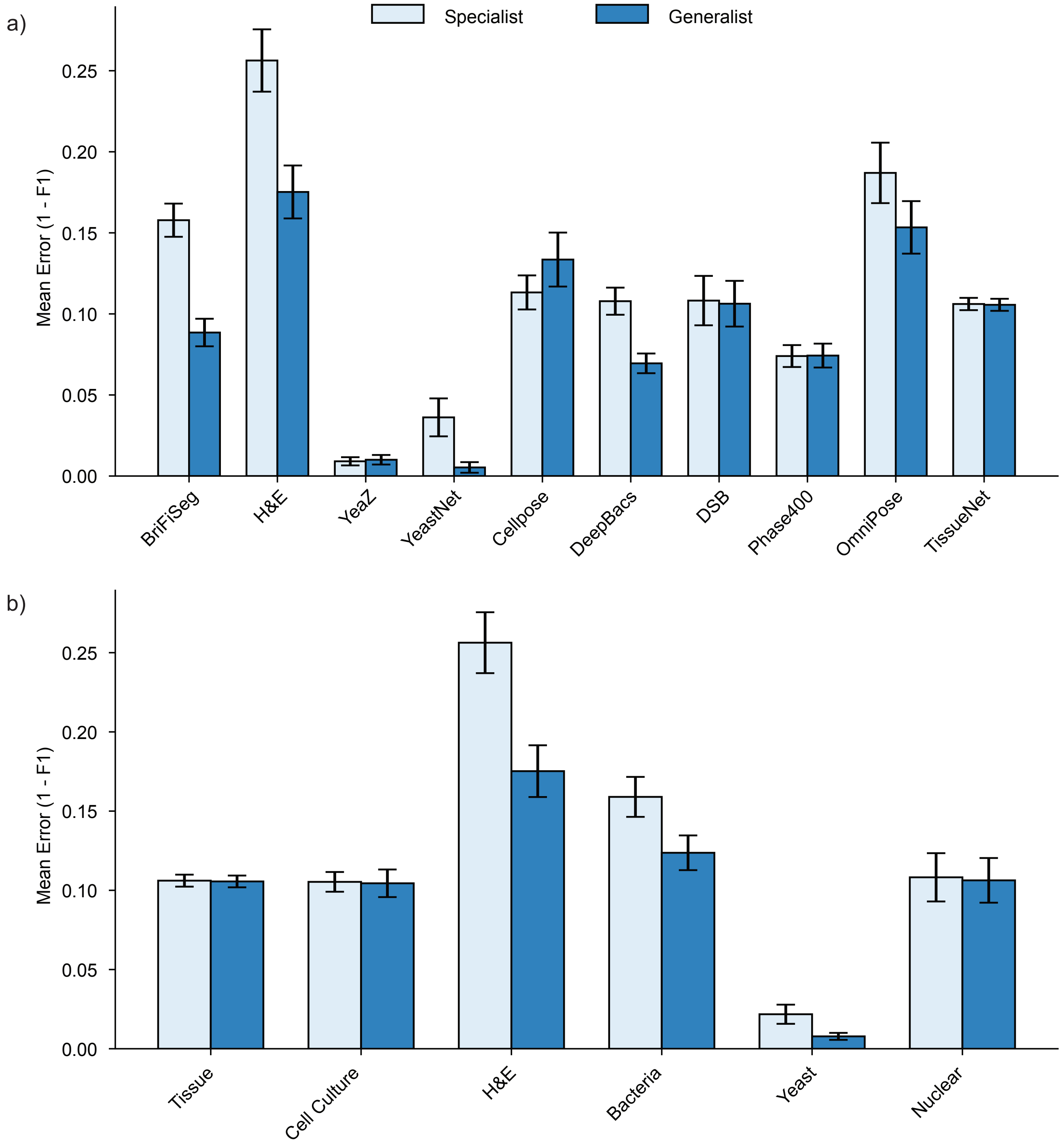


**Fig. S4**: **CellSAM-generalist vs CellSAM-specialist** We compare the performance of CellSAM when trained on individual datasets (specific) to when CellSAM is trained on all of the data (generalist). We see that CellSAM-generalist performs better across data types and datasets.

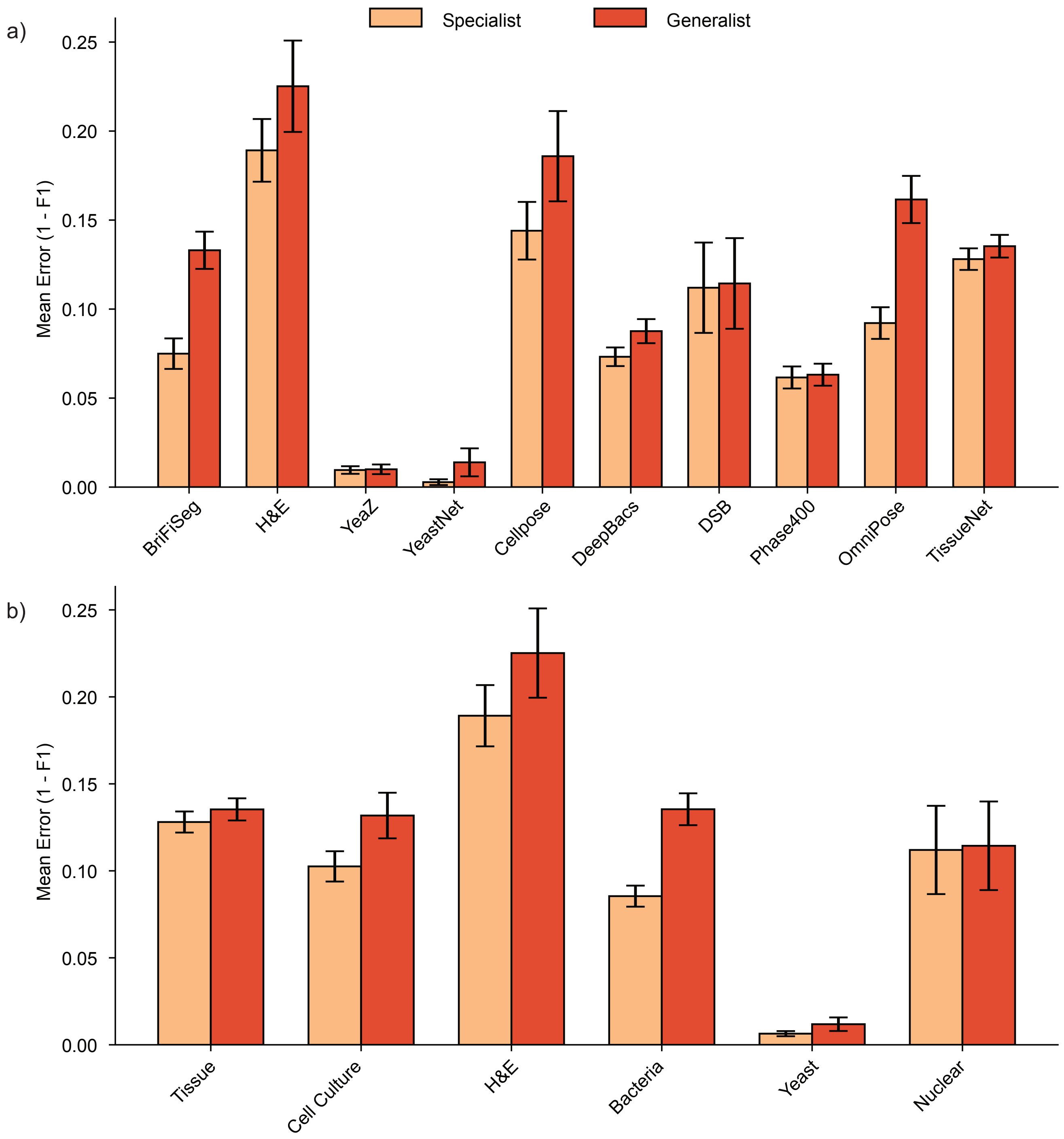


**Fig. S5**: **Cellpose-generalist vs Cellpose-specialist**. Here, we compare the performance of Cellpose when trained on individual datasets (specific) to when Cellpose is trained on all the data (generalist). Cellpose consistently performs better when trained on a specific dataset but does not perform well when trained across multiple datasets.

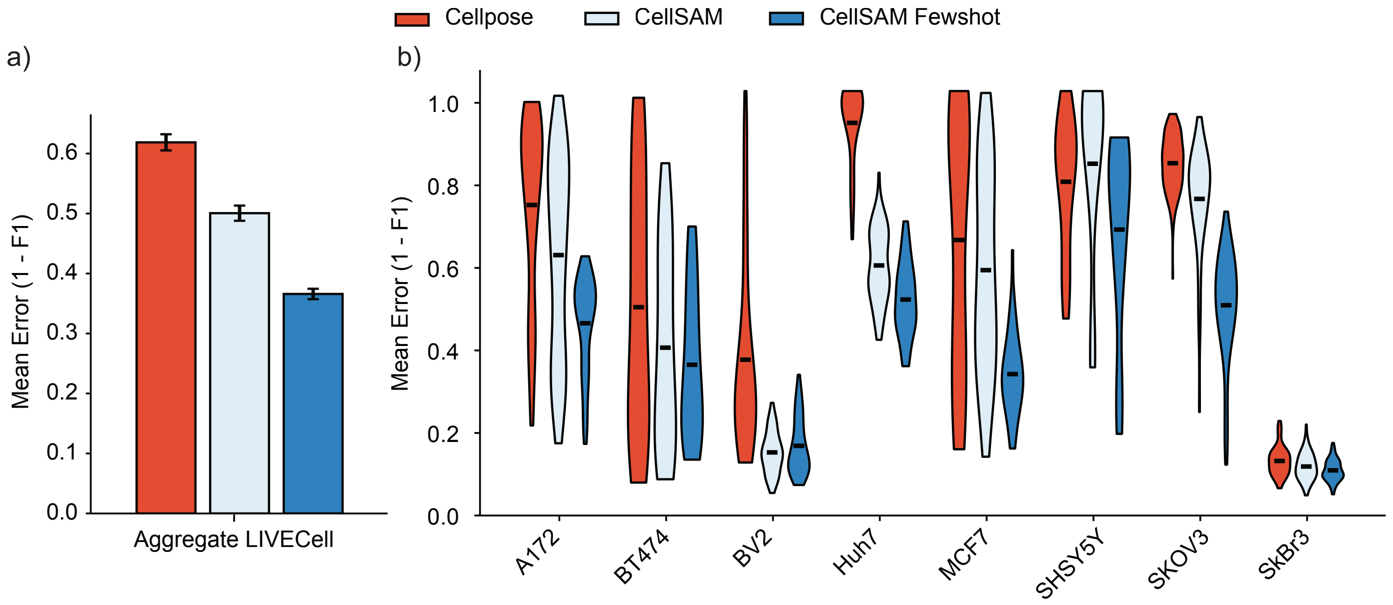


**Fig. S6**: a) Zero-shot performance of CellSAM-generalist and Cellpose-generalist on the LIVECell dataset. Here, we show significant improvement of CellSAM over Cellpose on an unseen dataset (from 0.13 to 0.40 in F1). b) CellSAM-generalist performance stratified by cell line. We analyzed both zero-shot and few-shot (10 samples per cell line) performance. We saw that few-shot improves CellSAM-generalist on LIVECell for most cell lines.

A172

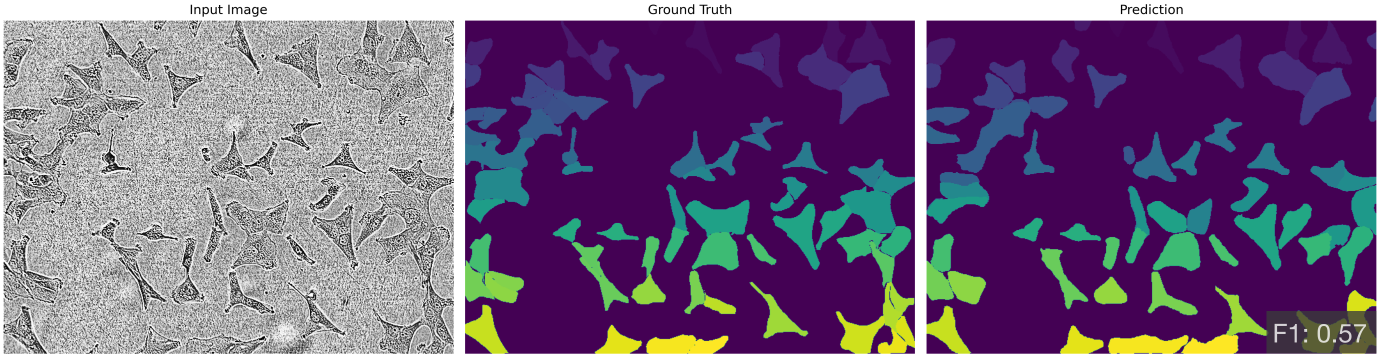


BT474

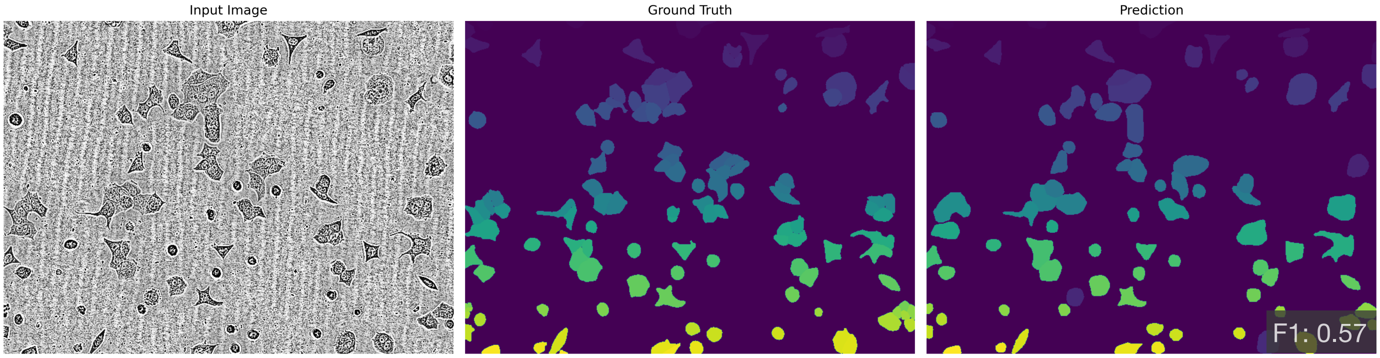


BV2

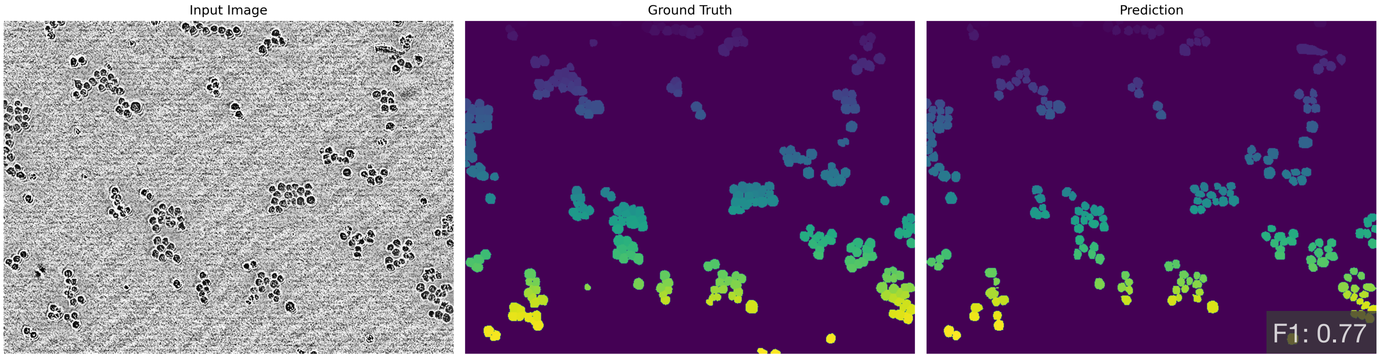


Huh7

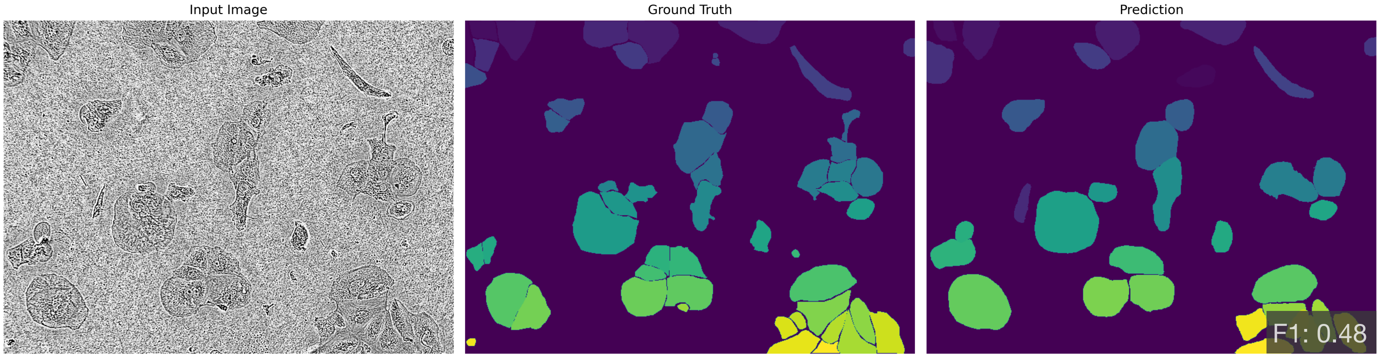

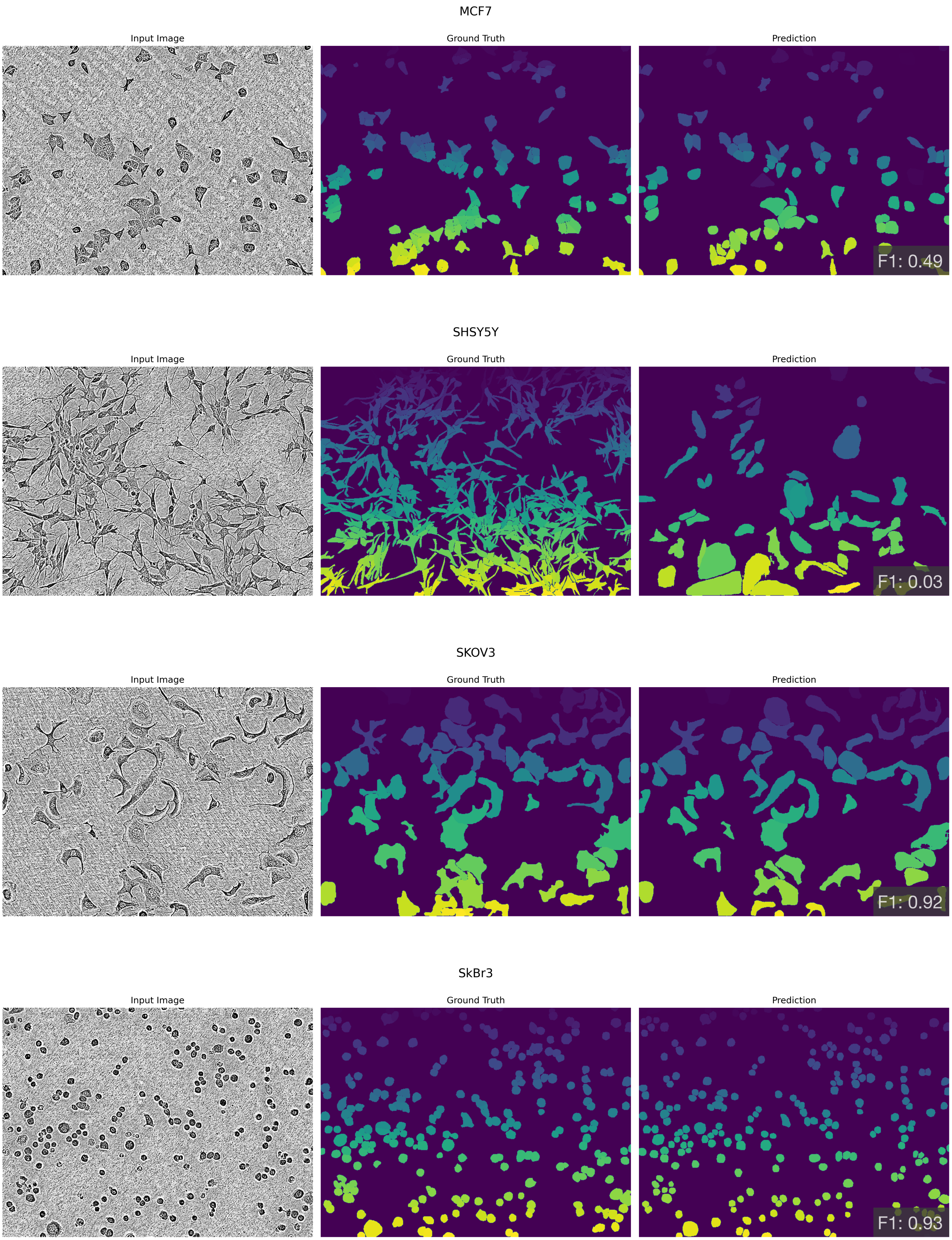


**Fig. S7**: Representative qualitative results of CellSAM-generalist 10-shot performance on the LIVECell dataset, one panel for each cell line.
